# Bidirectional Crosstalk Between Sleep and Pulmonary Arterial Hypertension

**DOI:** 10.1101/2025.07.23.666314

**Authors:** Seun Imani, Aymen Halouani, Jacob Dahlka, Nathan Burgess, Samar Antar, You-Yang Zhao, Stephen Y Chan, James D West, Matthew Weston, Yassine Sassi

## Abstract

**Background:** Pulmonary Arterial Hypertension (PAH) is a devastating cardiopulmonary disease characterized by pulmonary vascular remodeling due to vascular cells dysfunction. Among other clinical signs, emerging data suggest poor sleep quality in patients with PAH; however, how poor sleep impacts hemodynamic burden, symptom severity, and pulmonary vascular remodeling during PAH progression remains unknown.

**Methods:** We used two models of sleep disturbances (sleep fragmentation and chronic jet lag) and different mouse models of PAH to determine the effects of poor sleep on PAH. We carried out timepoint quantitative RT-PCR analyses to define the clock gene oscillations in human pulmonary artery smooth muscle cells (PASMCs) isolated from non-PAH and PAH patients. Bulk RNA-sequencing analysis, immunostaining, and proliferation assays were used to explore the mechanisms by which poor sleep impacts PAH. Electroencephalogram and Electromyogram recordings (EEG/EMG) were used to determine whether PAH affects sleep in mice.

**Results:** Poor sleep exacerbated right ventricular dysfunction, pulmonary vascular remodeling, and PAH. RNA-seq and immunostaining analyses showed that poor sleep induces lung inflammation. Inflammation affected the pulmonary vascular molecular clock to drive PASMC hyperproliferation and increased migration. SMC-specific deletion of Bmal1 protected mice from RV dysfunction, pulmonary vascular remodeling, and PAH. EEG/EMG measurements demonstrated that PAH causes poor sleep quality in mice. Lastly, we showed that improving sleep via melatonin delivery and blunting inflammation with clodronate inhibited pulmonary vascular remodeling and PAH.

**Conclusions:** Our study demonstrates that PAH causes poor sleep which in turn induces inflammation, increases PASMC proliferation, and exacerbates PAH. This suggests that the relationship between PAH and poor sleep is a self-amplifying cycle, and that a combination of hypnotic and anti-inflammatory drugs may give PAH patients better clinical outcomes.

**Clinical Perspective:** *What is New?:*

- PAH causes poor sleep quality which in turn induces lung inflammation, PASMC proliferation, and exacerbated PAH.
- Bmal1 rhythmicity is disrupted, and its expression is increased in lungs of patients with PAH.
- SMC-specific deletion of Bmal1 protects from PAH.
- Improving sleep quality and blunting inflammation attenuates PAH.

*What Are the Clinical Implications?:*

- Bmal1 represents a promising disease-modifying agent with potential for clinical translation in the treatment of PAH.
- A combination of hypnotic and anti-inflammatory drugs has therapeutic potential to treat PAH.

## INTRODUCTION

Pulmonary arterial hypertension (PAH) is a devastating cardiopulmonary disease characterized by pulmonary vascular remodeling that results from the dysfunction of pulmonary vascular cells, leading to right ventricular hypertrophy and right heart failure^1,2^. Despite some advancements in managing PAH, it remains a debilitating condition with a limited survival rate (with a median survival of 2.8 years from diagnosis)^3^. Treatment options only slightly alleviate symptoms, with a few modestly affecting pulmonary vascular remodeling^4,5^. Moreover, not all PAH patients respond clinically or hemodynamically to current treatments, highlighting the need to consider additional factors that contribute to PAH pathology. This emphasizes the importance of gaining insights into potential risk factors and developing more effective therapeutic interventions. Recent studies have reported poor sleep quality in patients with PAH^6–10^; however, the impact of sleep disturbances on hemodynamic parameters, symptom severity, and PAH progression has not been characterized. Therefore, it is imperative to explore the molecular connections between sleep disturbances and major pathological events that drive pulmonary vascular remodeling and the progression of PAH.

The timing of sleep is regulated by an internal molecular clock controlled by clock genes that oscillate nearly every 24 hours, forming the circadian rhythm^11,12^. This rhythm influences sleep-wake patterns and is essential for key physiological processes, including cell proliferation and blood pressure regulation^13,14^. The central (molecular) clock, located in the suprachiasmatic nucleus of the hypothalamus, synchronizes the molecular clocks found in peripheral tissues^15,16^; however, these peripheral clocks can also be influenced by other cues, like sleep, in a tissue-specific manner^17,18^. Sleep affects the rhythmic expression of genes and proteins vital for lung homeostasis, and disturbances in sleep may disrupt the circadian oscillations of the pulmonary molecular clock, potentially leading to pulmonary diseases.

Basic Helix-Loop-Helix ARNT Like 1 (BMAL1) and Clock Circadian Regulator (CLOCK), which regulate downstream clock genes and drive the molecular clock machinery^19^, form a heterodimer complex (BMAL1:CLOCK) that controls the rhythmic expression of most genes, including clock and cell cycle genes^20–22^. Disruptions in the oscillations and expressions of these clock genes have been associated with various diseases^23–26^. Sleep disruptors, such as jet lag and sleep fragmentation, have been linked to the risk and progression of different human diseases^27–29^; yet, the existing literature provides limited insight into the relationship between sleep/circadian clock disruptions and PAH.

In this study, we employed two models of sleep disturbance (sleep fragmentation and chronic jet lag), various mouse models of PAH, and in vitro methods to investigate the impact of poor sleep on PAH. We examined the circadian rhythmicity and expression of clock genes in primary vascular cells and lungs from patients with PAH. Using electroencephalogram and electromyogram measurements, we investigated the effect of PAH on sleep quality in mice. We explored the therapeutic potential of targeting Bmal1 in treating PAH, as well as the benefits of improving sleep quality and blunting lung inflammation on pulmonary vascular remodeling and PAH.

## METHODS

The data that support the findings of this study are available from the corresponding author on reasonable request. An expanded Methods section is provided in the Supplemental Material.

### Animal studies

Wild-type mice were obtained from Envigo. Age- and sex-matched mice were used for all in vivo studies. All mice were group-housed under a 12-hour light and 12-hour dark cycle (unless in the Chronic Jet Lag experiments, where the light/dark cycle was advanced by 6 hours every 2 days). Mice had ad libitum access to food and water. All animal protocols were approved by the VT animal protocol under IACUC guidelines. Lung-specific IL6 overexpression transgenic mice (Clara10-IL6 Tg+) were kindly provided by Dr. Aaron Waxman^30^. The generation of smooth muscle-specific doxycycline-inducible BMPR2 mutant mice (SM22-rtTAxTet-BMPR2^R899X^) was previously described^31^. The generation of mice with endothelial cell-specific deletion of Egln1 (Tie2Cre;Egln1^fl/fl^) was previously described^32^. To generate smooth muscle cell-specific Bmal1 knockout (SMC-Bmal1 KO) and endothelial cell-specific knockout (EC-Bmal1 KO) mice, we crossed Bmal1^flox/flox^ mice (JAX strain #007668) with tamoxifen-inducible Myh11-CreER^T2^ (obtained from Dr. Gary Owens^33^) and Cdh5-CreER^T2^ (originally produced in Dr. Ralf Adams lab at The Max Planck Institute for Molecular Biomedicine and received from Dr. Gary Owens) mice, respectively. Homozygous littermates Bmal1^flox/flox^ but lacking Cre were used as controls. To activate Bmal1 deletion, 8-10-week-old mice received daily intraperitoneal injection of Tamoxifen (40mg/Kg, Sigma) for 2 consecutive weeks.

### Human Samples

Human lungs, human pulmonary artery smooth muscle cells and human pulmonary artery endothelial cells were obtained from the Pulmonary Hypertension Breakthrough Initiative (PHBI). Patient enrollment and the standardized tissue-processing protocol for the PHBI have been described previously^34,35^. Patient characteristics are described in Tables S1-S3.

## RESULTS

### Sleep fragmentation exacerbates PAH

To investigate the effect of sleep disturbances on PAH, we subjected wild-type mice to a model of chronic sleep fragmentation (SF) by placing them in a SF chamber with a bar that sweeps the bottom of the cage every 2 minutes during the light cycle. Mice placed in the SF chamber with a stationary bar were used as controls. Mice were then exposed to the Sugen/Hypoxia (Su/Hx) model of PAH (Figure 1A), and cardiac hemodynamic measurements were performed 4 weeks later. Mice subjected to SF had significantly increased right ventricular systolic pressure (RVSP) (Figure 1B) and cardiomyocyte hypertrophy, as shown by increased N-terminal pro B-type natriuretic peptide (NT-proBNP) mRNA level and cardiomyocyte size (Figure 1C-D). Next, we evaluated the impact of SF on pulmonary vascular remodeling by performing Hematoxylin and Eosin staining on mice lung sections from the different groups. Mice under SF showed significantly increased pulmonary vascular remodeling, as indicated by increased pulmonary arterial medial thickness (Figure 1E).

**Figure 1:**
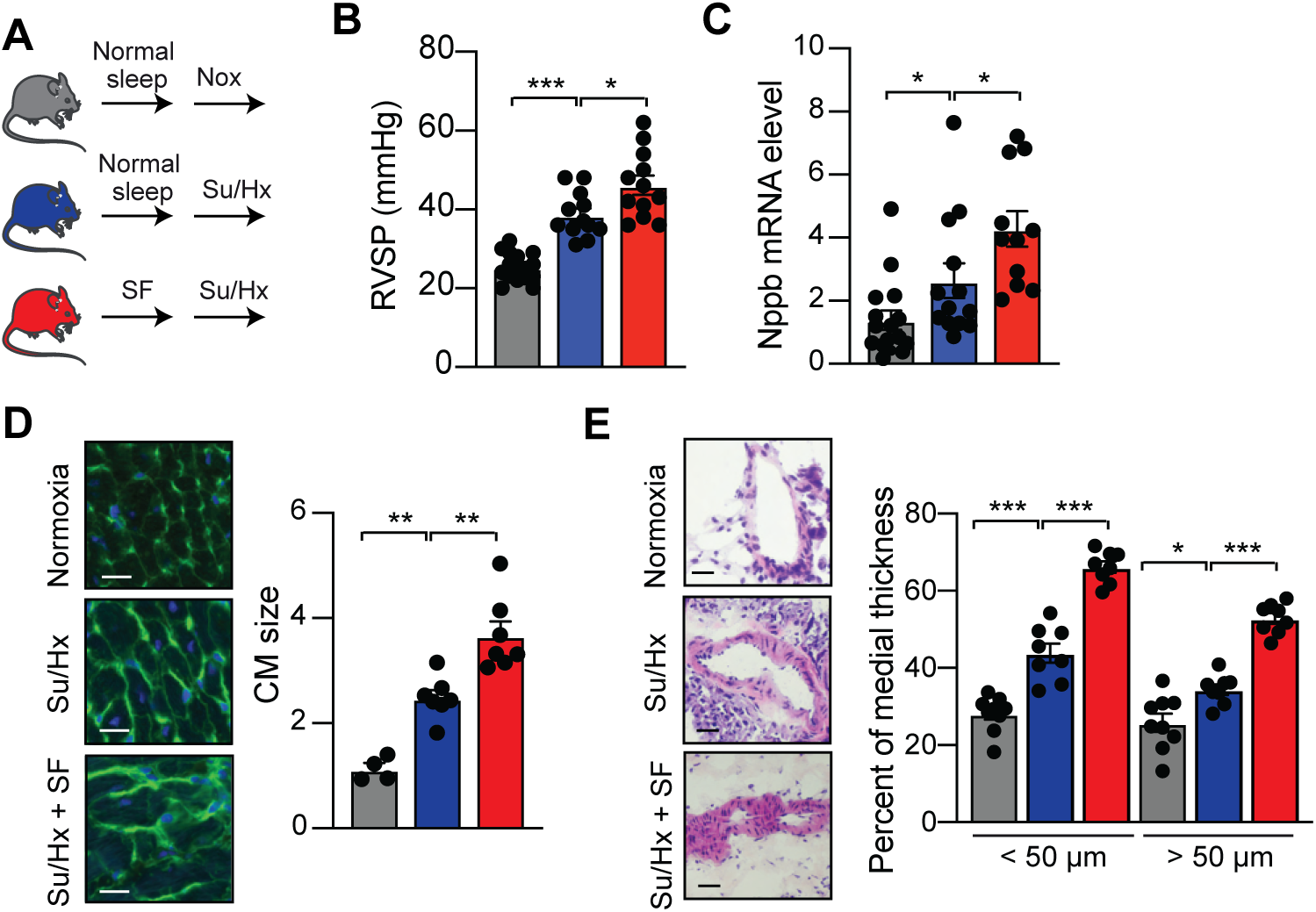
Sleep Fragmentation exacerbates PAH. (**A**) Experimental design. WT mice were subjected to 16 weeks of SF followed by exposure to a Sugen/hypoxia model of PAH. (**B**) RVSP of the indicated groups. (**C**) Natriuretic Peptide B (Nppb) mRNA level in the hearts of the indicated groups. (**D**) (Left) Representative WGA-stained RV sections. Scale bar: 50 μm. (Right) Quantitative analysis. (**E**) (Left) Representative H&E-stained pulmonary arteries sections from the indicated groups. Scale bar: 50 μm. (Right) Percentage of arteries medial thickness. n= 12-15 mice/group. Data were analyzed by 1-way ANOVA. *P<0.05, ** P<0.01, ***P<0.001.

To determine whether the increased PAH phenotype is due to stress induced by the moving sweep bar, mice were placed in SF chambers with the sweep bar set to move along the bottom of the cage every 2 minutes during the night cycle (when the mice are active). Control mice were placed in SF chambers with a stationary bar. Nighttime sweeping of the bar did not affect cardiac and pulmonary vascular remodeling (Figure S1), indicating that PAH exacerbation is not due to physical or psychological stress from the used model. These results suggest that SF exacerbates PAH.

### Poor sleep worsens PAH in genetically predisposed PAH mice

Next, to test whether poor sleep exacerbates PAH in a separate genetic model of this disease, we characterized the impact of SF on PAH using mice with endothelial cell-specific deletion of Egln1 (Tie2Cre;Egln1^fl/fl^) that spontaneously develop severe PAH^32^. Given that the homozygous Egnl1^-/-^ mice die within 3 months after birth (Figure 2A), we used heterozygous Egnl1^+/-^ mice, which are viable and spontaneously develop PAH. SF exacerbated PAH in Egnl1^+/-^ mice with elevated RVSP and RV hypertrophy (both at the tissue and cellular levels), and increased pulmonary vascular remodeling (Figure 2B-F). These results suggest that SF worsens PAH in genetically predisposed mice.

**Figure 2:**
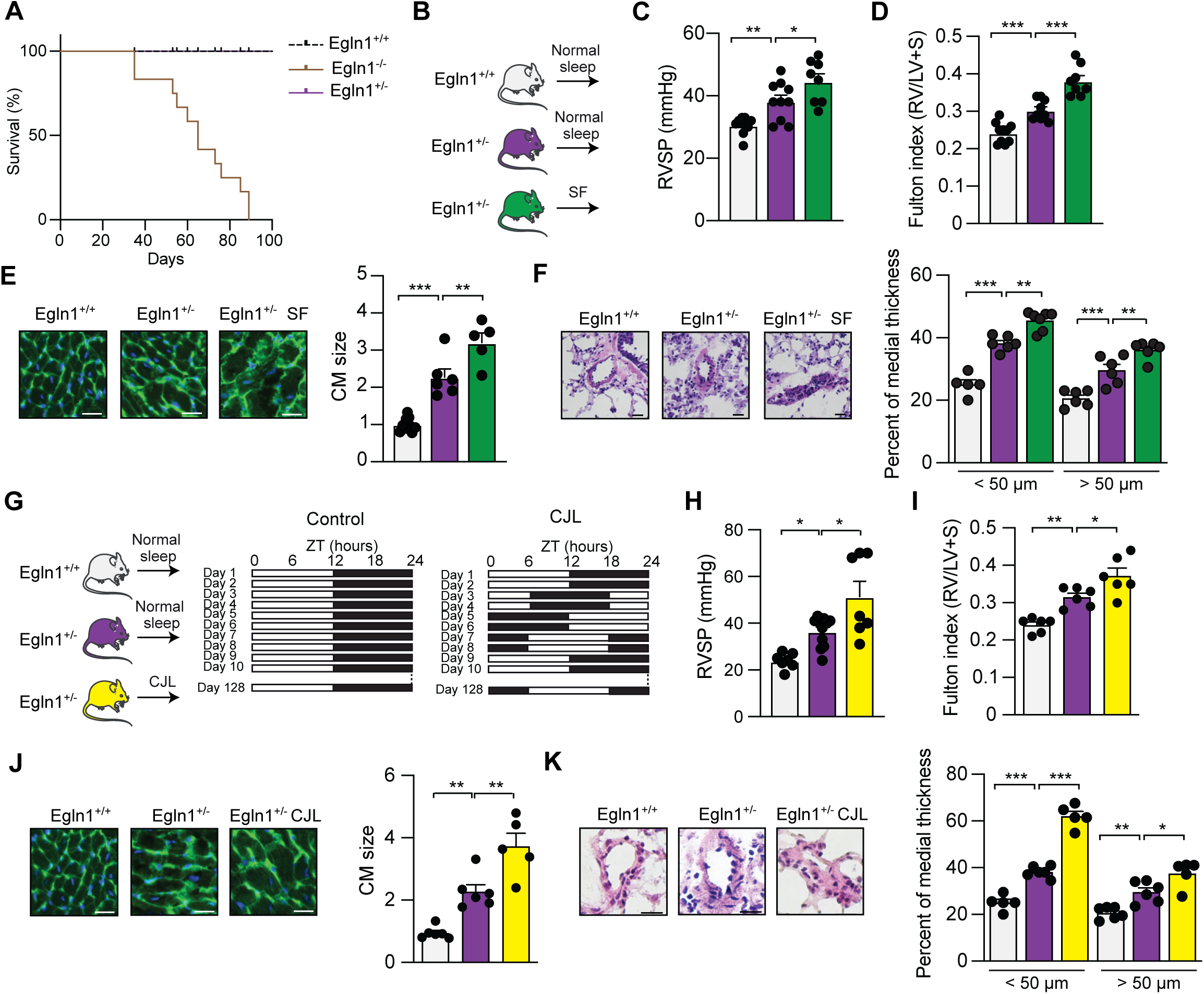
Poor sleep exacerbates PAH in genetically predisposed mice. (**A**) Kaplan Meier survival analysis of homozy-hous and heterozygous Egln1 KO mice and their WT littermates. (**B**) Experimental design of the SF study. (**C**) RVSP and (**D**) Fulton index of the indicated groups. (**E**) (Left) Representative WGA-stained RV sections. Scale bar: 50 μm. (Right) Quantita-tive analysis. (**F**) (Left) Representative H&E-stained pulmonary arteries sections from the indicated groups. Scale bar: 50 μm. (Right) Percentage of arteries medial thickness. n= 5-10 mice/group. (**G**) Experimental design and light regime for chronic jet lag treatment. Black bars represent dark phase and hollow bars represent light phase. (**H**) RVSP and (**I**) Fulton index of the indicated groups. (**J**) (Left) Representative WGA-stained RV sections. Scale bar: 50 μm. (Right) Quantitative analysis. (**K**) (Left) Representative H&E-stained pulmonary arteries sections from the indicated groups. Scale bar: 50 μm. (Right) Percent-age of arteries medial thickness. n= 5-7 mice/group. Data were analyzed by 1-way ANOVA.

To exclude that poor sleep-exacerbated PAH is solely due to the used SF model, we used the chronic jet lag (CJL) model of sleep disturbance. In this model, mice were placed in a circadian cabinet and were subjected to circadian disruption through phase advances of 6 hours every 2 days. Quantitative PCR analyses, performed on RNA extracted from the lungs of control and CJL mice that were sacrificed every 4 hours over a 24 hour period, revealed robust circadian oscillations of core clock genes in control mice, and disrupted clock oscillations in the lungs of CJL-exposed mice (Figure S2). Next, we subjected Egnl1^+/-^ mice to CJL (Figure 2G). Egnl1^+/-^ mice and littermate controls under constant 12:12-hours light/dark cycle were used as controls (Figure 2G). CJL significantly worsened PAH in Egnl1^+/-^ mice, as shown by increased RVSP (Figure 2H), cardiac hypertrophy at the tissue and cellular levels (Figure 2I-J), and pulmonary vascular remodeling (Figure 2K). Collectively, these findings demonstrate that poor sleep is a significant risk factor for PAH progression.

### Poor sleep induces lung inflammation

To determine the mechanisms underlying poor sleep-mediated exacerbation of PAH, lungs from Su/Hx mice with or without SF were harvested (under the same circadian time), and bulk RNA sequencing was performed to reveal transcriptomic enrichment analyses. Among the regulated mRNAs, 285 downregulated and 560 upregulated genes were differentially expressed upon poor sleep induction. To identify molecular functions of the mRNAs altered by sleep disturbance, Gene Ontology (GO) and Kyoto Encyclopedia of Genes and Genomes (KEGG) pathway enrichment analyses were then performed for the identified mRNAs. The functional categories of genes significantly enriched upon sleep disturbance using GO and KEGG pathway analyses are summarized in Figure 3A-B. The majority of regulated pathways in lungs of SF mice were associated with circadian rhythm and inflammation (Figure 3B). With these enrichment analyses, we hypothesized that poor sleep induces immune cells activation (and recruitment), thereby causing macrophage accumulation in the lung. To test this hypothesis and independently validate the findings of changes in inflammation-associated pathways, we antibody-stained tissue cryosections of lungs obtained from control, Su/Hx and Su/Hx mice subjected to SF for the macrophage marker CD68 (Figure 3C). Quantitative analysis confirmed a significant increase of CD68 positive macrophages in the lungs of sleep disturbed mice (Figure 3C). These results indicate that poor sleep exacerbates the inflammatory response in the lungs of PAH-diseased mice.

**Figure 3:**
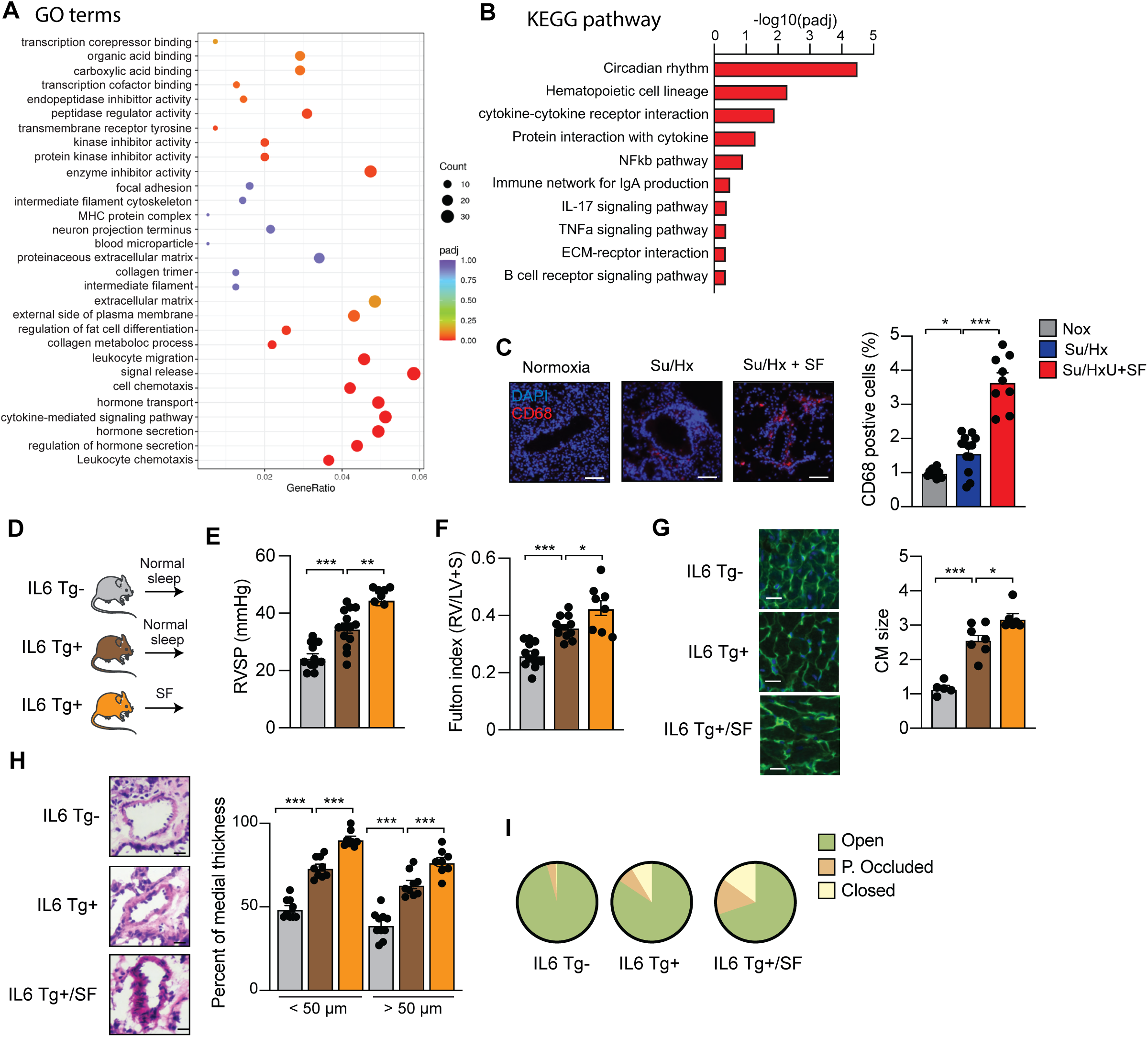
Poor sleep triggers inflammation in PAH-diseased mice. (**A**) Scatter plot of top 30 gene ontology (GO) terms enriched by differentially expressed genes (DEGs) in Su/Hx+SF vs Su/Hx lungs (n=4 mice/group). ‘padj’ is the P-value adjusted using the Benjamini-Hochberg procedure. ‘Count’ is the number of genes enriched in a GO term. ‘GeneRatio’ is the percentage of total DEGs in the given GO term. (**B**) KEGG pathway enrichment analysis.The significant pathway for differentially expressed genes in Su/Hx+SF vs Su/Hx lungs. (**C**) CD68+ cells (red) in the respective lungs with quantification. Scale bar: 20 µm. (**D**) Experimental design of the study. (**E**) Right ventricular systolic pressure (RVSP) of the indicated groups. (**F**) Fulton index of the indicated groups. (**G**) (Left) Representative WGA-stained RV sections to assess hypertrophy of cardiac myocytes. Scale bar: 50 μm. (Right) Quantitative analysis. (**H**) (Left) Representative H&E-stained pulmonary arteries sections from the indicated groups. Scale bar: 25 µm. (Right) Percentage of arteries medial thickness. (**I**) Percentage of occluded pulmonary arteries of the indicated groups. n= 10-15 mice/group. * P<0.05, ** P<0.01, ***P<0.001. Data were analyzed by 1-way ANOVA.

To explore the effects of cytokines and poor sleep on PAH, we used mice with lung-specific Interleukin-6 overexpression (Clara10-IL6 Tg+) and subjected them to SF or normal sleep (Figure 3D). The WT littermates (IL6 Tg-) were placed in SF chamber with a stationary bar and used as controls. Mice with Interleukin-6 overexpression spontaneously developed PAH and poor sleep aggravated their phenotype. IL6 Tg+ mice displayed all the hallmarks of PAH (i.e., increased RVSP and Fulton index) and sleep fragmentation induced a marked increase in these parameters (Figure 3E-F). Furthermore, poor sleep exacerbated cardiomyocyte hypertrophy and increased pulmonary arterial medial thickness in the IL6 Tg+ mice, as determined by histological analysis (Figure 3H). In addition, we found increased pulmonary arterial occlusion in IL6 Tg+ mice under SF protocol when compared to age-matched IL6 Tg+ mice without SF (Figure 3I). Bulk RNA-seq analysis on lungs isolated from IL6 Tg+ subjected or not to SF, confirmed the changes in inflammation-associated pathways in the lungs of mice subjected to SF (Figure S3). These results demonstrate that poor sleep induces lung inflammation in PAH-diseased mice.

### The principal drivers of the molecular clock control PASMCs proliferation and are disrupted in PASMCs from patients with PAH

The key events leading to pulmonary vascular remodeling in PAH are typified by hyperproliferation of the pulmonary artery endothelial (PAECs) and smooth muscle cells (PASMCs), suggestive of dysfunction in the cell cycle dynamics; which are gated by the molecular clock^36,37^. Thus, we explored the role of clock genes in PAH. We first defined the presence of circadian clock oscillations in PAECs and PASMCs isolated from healthy donors. Cells were synchronized with dexamethasone, and samples were collected every 4 hours over 24 hours to assess the mRNA levels of clock genes. Robust circadian oscillations of Bmal1 and Clock (the major components of the circadian clock) were detected in PASMCs, and lesser oscillations in PAECs (Figure 4A). This finding suggests that circadian regulation of pulmonary vascular cell function is mainly driven by PASMCs. We then investigated clock oscillations in PASMCs isolated from patients with PAH and found that Bmal1 and Clock oscillations are disrupted in PAH-diseased PASMCs compared to healthy PASMCs, suggesting disruption of the molecular clock in PAH (Figure 4B). We next quantified pulmonary Bmal1 and Clock mRNA levels in lungs from non-PAH patients and from patients with idiopathic PAH. The PCR analyses revealed no significant changes in Bmal1 and Clock mRNA levels in diseased human lungs (Figure 4C). We next investigated the expression levels of Bmal1 and Clock in PASMCs isolated from non-PAH patients and from patients with idiopathic PAH. We found Bmal1 and Clock levels to be substantially higher in PASMCs from patients with clinical PAH than in healthy PASMCs (Figure 4D). Increase in Bmal1 and Clock expression disrupted the timed rhythms of these genes, causing flattened and abnormal amplitudes of their oscillations. These results indicate that Bmal1 and Clock levels are increased and their rhythmicity disrupted in PAH-diseased PASMCs.

**Figure 4:**
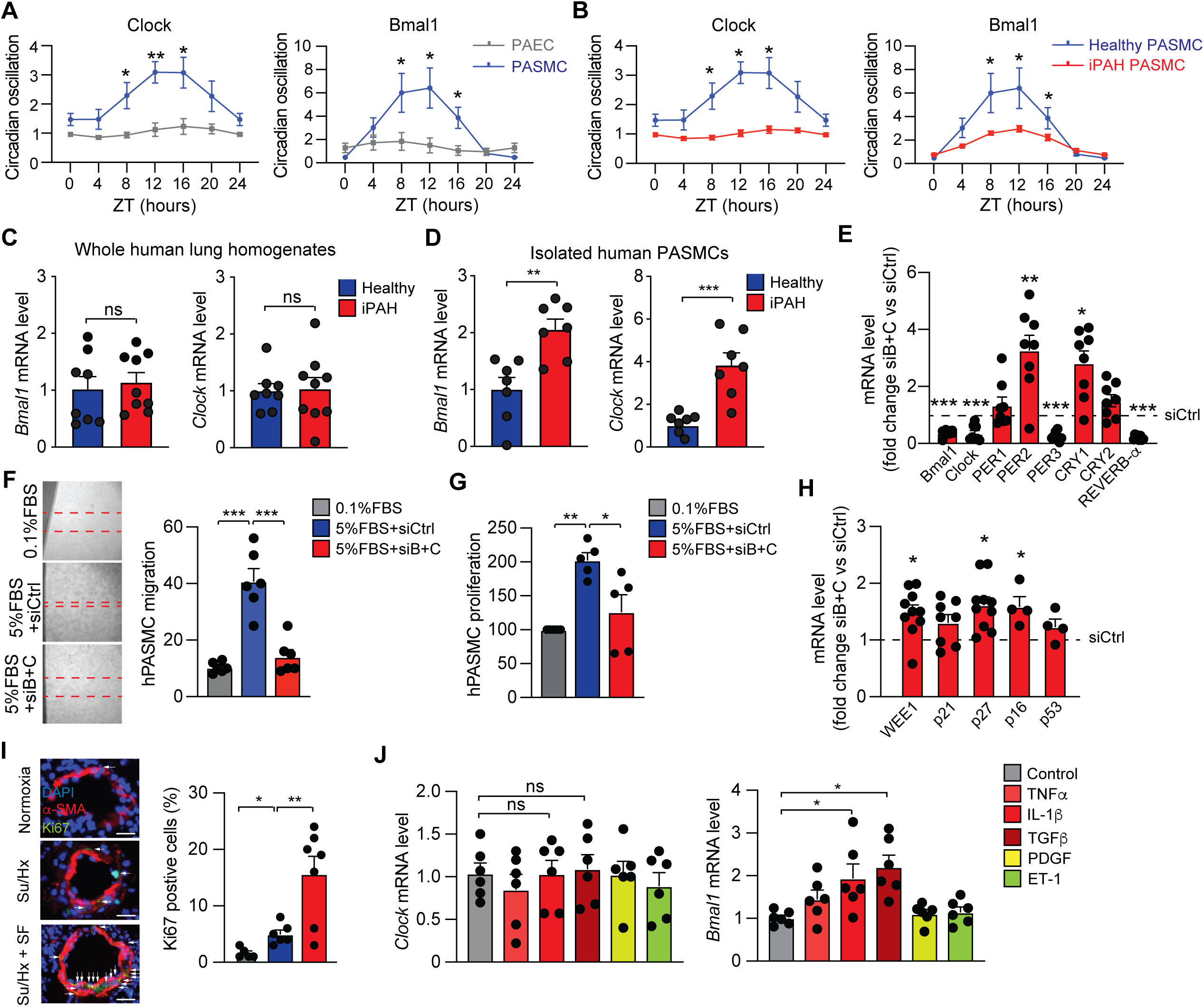
PAH affects circadian genes expression and rhythmicity. **(A)** Circadian Bmal1 and Clock oscillations in healthy PAECs and PASMCs. n= 6 patients/group. (**B**) Circadian Bmal1 and Clock oscillations in PASMCs from iPAH patients. n= 6 patients/group. **(C**) Bmal1 and clock mRNA levels in whole lung homogenates from healthy donors and patients with iPAH (n=8-9 patients/group). (**D**) Bmal1 and clock mRNA levels in isolated PASMCs from healthy donors and patients with iPAH (n=7 patients/group). **(E)** Circadian genes mRNA levels in PASMCs transfected with siCtrl or siBmal1+siClock (siB+C). n = 8 experiments performed in triplicate. (**F**) Migration and (**G**) proliferation of PASMCs in the presence of the indicated treatments. n = 5-6 experiments. (**H**) Cell cycle genes mRNA levels in PASMCs transfected with siCtrl or siBmal1+siClock. n = 4-10 experiments performed in duplicate. (**I**) Left) Representative lung sections stained with Dapi (blue), α-SMA (green) and Ki-67 (red). Scale bar: 50 μm. (Right) Quantitative analysis. n= 5-7 mice/group. (**J**) Bmal1 and clock mRNA levels in human PASMCs treated for 16 hours with TNFα (10 ng/mL), IL-1β (10 ng/mL), TGFβ (5 ng/mL), PDGF (10 ng/mL) or ET-1 (1 nM). n = 5-6 experiments performed in duplicate. * P<0.05, ** P<0.01, ***P<0.001. Data were analyzed by two-tailed t-test or 1-way ANOVA.

We next investigated whether increased Bmal1 and Clock expression affects PASMCs proliferation and migration. Knockdown of Bmal1 and Clock, using specific siRNAs, altered the expression of other clock genes further supporting that Bmal1 and Clock are key drivers of the molecular clock (Figure 4E). Knockdown of Bmal1/Clock suppressed the proliferation and migration of PASMC (Figure 4F-G). Additionally, quantitative PCR analyses revealed a significant increase in mRNA levels of cell cycle inhibitors P27, P16, and the G2/M checkpoint kinase WEE1 upon Bmal1/Clock knockdown (Figure 4H), suggesting that clock genes may regulate PASMC proliferation through G1/S and G2/M checkpoint control.

Next, we examined whether poor sleep-induced circadian genes regulation induces proliferation of PAMSCs in mice. Immunostaining analysis for a proliferating marker (ki-67) was performed on lung sections of Su/Hx mice with or without SF. The medial layer (PASMCs) of the pulmonary arteries of SF-exposed mice showed an increased level of Ki67-positive cells (Figure 4I), indicating that poor sleep induces aberrant proliferation of PASMCs to worsen pulmonary vascular remodeling and PAH.

Since poor sleep induces lung inflammation, we speculated that cytokines, released from inflammatory cells during poor sleep, may affect PASMC function by regulating the expression of clock genes to drive hyperproliferation of PASMCs and consequently increased pulmonary vascular remodeling. Therefore, we quantified Clock and Bmal1 expression in PASMCs treated with pro-PAH factors (i.e., cytokines, growth factor, and hormone). Our analysis revealed a significant upregulation of Bmal1 (but not Clock) in response to cytokines (Figure 4J). These results suggest that inflammation alters Bmal1 expression in PASMCs.

### Smooth Muscle cell-specific deletion of Bmal1 protects mice against PAH

Given that poor sleep triggers lung inflammation and that cytokines activate Bmal1 in PASMCs, we next assessed whether a SMC-specific deletion of Bmal1 protects mice from PAH. SMC-specific Bmal1 knockout mice (Myh11-CreER^T2^;Bmal1^fl/fl^ mice) were generated by crossing Bmal1^fl/fl^ mice with Myh11-CreER^T2^ mice. Myh11-CreER^T2^-Bmal1^fl/fl^ mice and their control littermates intraperitoneally received tamoxifen (40 mg/kg) for two consecutive weeks, and were then weekly injected with Sugen (20 mg/kg) for two weeks (i.e. 3 injections) and were maintained under hypoxic conditions (10% O_2_) for 4 weeks (Figure 5A). Hemodynamic, histological, and morphometric measurements revealed that SMC-specific deletion of Bmal1 protects mice against PAH. Control mice developed hallmark features of PAH, including elevated RVSP and Fulton index (Figure 5B-C). In contrast, Bmal1-deficient mice exhibited a marked reduction in these parameters, indicating a protective effect of Bmal1 deletion against PAH development (Figure 5B-C). Furthermore, histological analyses showed that Bmal1 deletion prevented RV cardiomyocyte hypertrophy (Figure 5D) and significantly reduced the percentage of pulmonary arterial wall thickness (Figure 5E). Moreover, lungs of SMC-specific Bmal1 deficient mice displayed a significant reduction of proliferating PASMCs (Figure 5F). We next generated mice with an endothelial cell-specific deletion of Bmal1 (Cdh5-CreER^T2^-Bmal1^fl/fl^ mice) by crossing Bmal1^fl/fl^ mice with Cdh5-CreER^T2^ mice. Endothelial cell-specific Bmal1 deficient mice did not show any phenotype at basal level (normoxic condition) and were not protected against PAH (Figure S4). These results indicate that a smooth muscle cell-specific deletion of Bmal1 protects mice from PAH.

**Figure 5:**
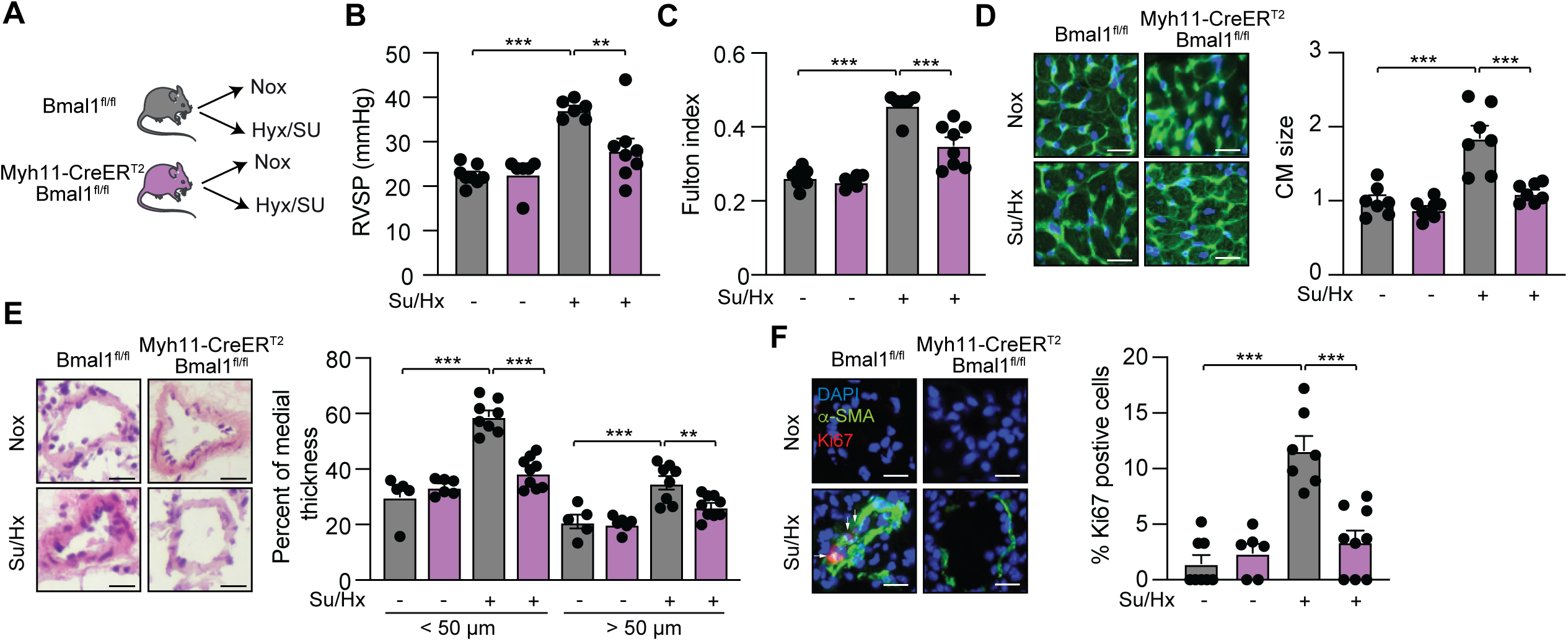
SMC-specific Bmal1 deletion protects mice against PAH. (A) Design of the study. (B) RVSP of the indicated groups. (C) Fulton index of the indicated groups. (D) (Left) Representative WGA-stained RV sections. Scale bar: 50 μm. (Right) Quantitative analysis. (E) (Left) Representative H&E-stained pulmonary arteries sections from the indicated groups. Scale bar: 50 μm. (Right) Percentage of arteries medial thickness. (F) (Left) Representative lung sections stained with Dapi (blue), α-SMA (green) and Ki-67 (red). Scale bar: 50 μ m. (Right) Quantitative analysis. n= 7-9 mice/group. Data were analyzed by 2-way ANOVA.

### BMAL1 is upregulated in the medial layer of remodeled vessels in human PAH

The analysis of C-C Chemokine Receptor 2 (CCR2) expression by immunofluorescence showed an increased number of CCR2 positive cells in the lungs of patients with PAH (Figure 6A), suggesting a high infiltration of circulating monocytes and an enhanced monocyte-to-macrophage differentiation chronically in PAH. The increased number of inflammatory cells in the lungs of patients with PAH prompted us to quantify the cytokines levels in the lungs of patients with PAH. Immunofluorescence analyses revealed a significant increase of IL-6 level in the lungs of patients with PAH (Figure 6B). BMAL1 signal was mainly detected in pulmonary vessel cells and co-localized with ɑ-smooth muscle actin (α-SMA) in the medial layer of the pulmonary arterial walls of PAH patients in comparison with the control vessels of non-PAH lungs (Figure 6C). The upregulation of Bmal1 level in the lungs of patients with PAH and the protective effects of Bmal1 deletion in SMC indicate that BMAL1 inhibition represents a potential disease-modifying approach in the treatment of PAH.

**Figure 6:**
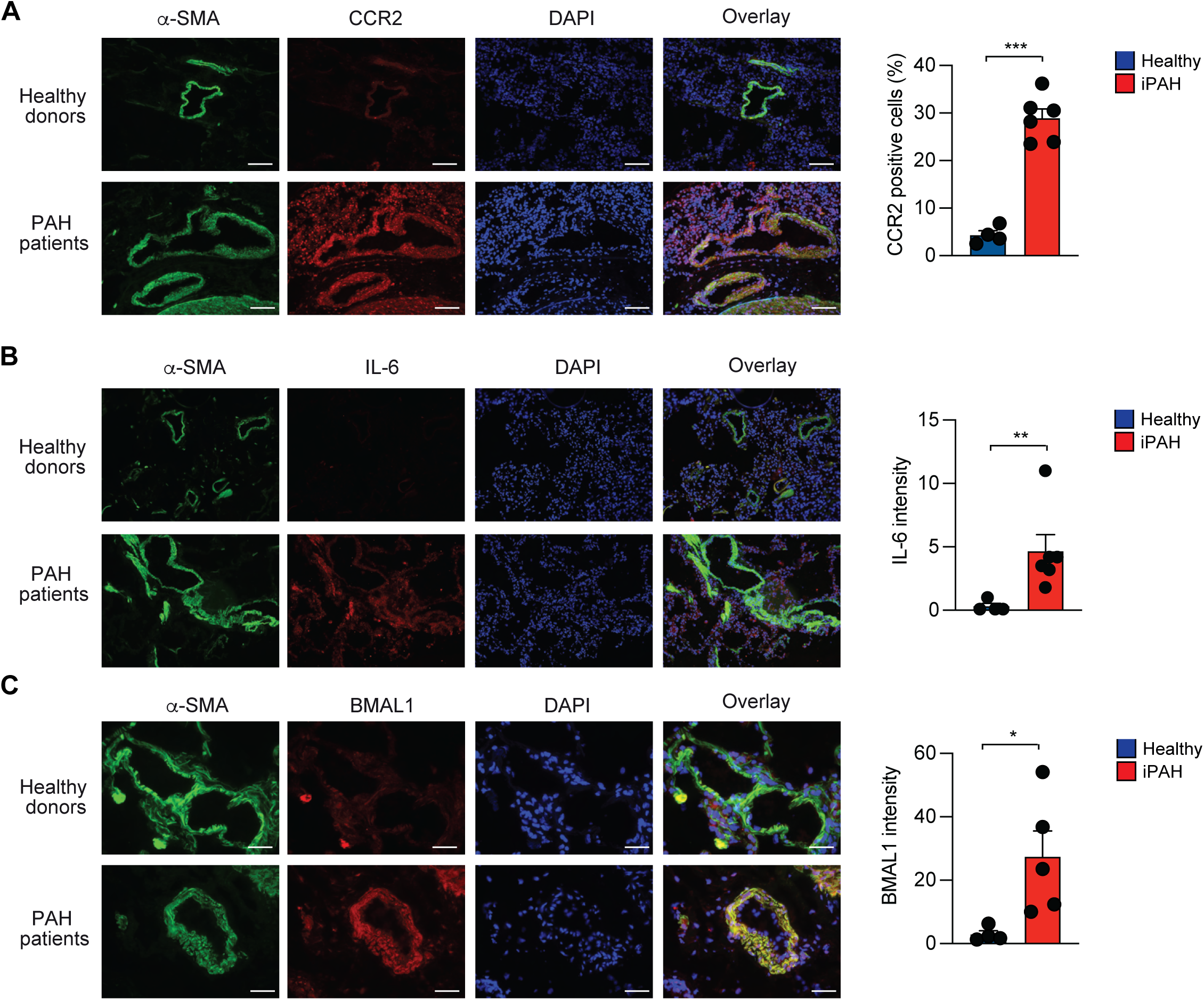
Increased macrophages infiltration, cytokines production and Bmal1 expression in lungs of patients with PAH. (**A**) (Left) Representative lung sections stained with Dapi (blue), α-SMA (green) and CCR2 (red). Scale bar: 20 μm. (Right) Quantitative analy-sis. n= 4-6 patients/group. (**B**) (Left) Representative lung sections stained with Dapi (blue), α-SMA (green) and IL-6 (red). Scale bar: 20 μ m. (Right) Quantitative analysis. n= 4-6 patients/group. (**C**) (Left) Representative lung sections stained with Dapi (blue), α-SMA (green) and BMAL1 (red). Scale bar: 20 μm. (Right) Quantitative analysis. n= 4-6 patients/group. * P<0.05, ** P<0.01, ***P<0.001. Data were analyzed by two-tailed t-test.

### PAH causes poor sleep in mice

Since emerging data suggest that patients with PAH experience poor sleep, we examined whether PAH causes poor sleep in mice. We used wireless electroencephalography (EEG) and electromyography (EMG) recording electrodes to monitor sleep and wakefulness in mice. To validate the effectiveness of the EEG/EMG system for assessing sleep quality in mice, we implanted EEG/EMG electrodes in WT mice and subjected them to SF or normal sleep. The sleep and wake states were measured continuously for eight weeks. Mice subjected to SF had a significantly higher wake count (Figure S5A), decreased total sleep time (Figure S5B) and Non-Rapid Eye Movement (NREM) sleep during the light cycle (Figure S5C), indicating that EEG/EMG telemetry is a powerful tool to assess sleep quality in mice. Next, we recorded EEG signals from the brain and EMG signals from muscles to quantify sleep and wake states and transitions in Su/Hx or normoxic mice (Figure 7A). Remarkably, PAH increased the wake counts during the rest period (Figure 7B) and decreased sleep and NREM counts during the same period (Figure 7C-D). Furthermore, the hypnogram data showed that Su/Hx mice had fewer transitions between NREM and REM sleep and spent most of their time awake when compared to healthy mice (Figure 7E-F). The percentage of time spent in sleep and NREM was significantly reduced in Su/Hx mice (Figure 7F). These results indicate that Su/Hx causes poor sleep in mice.

**Figure 7:**
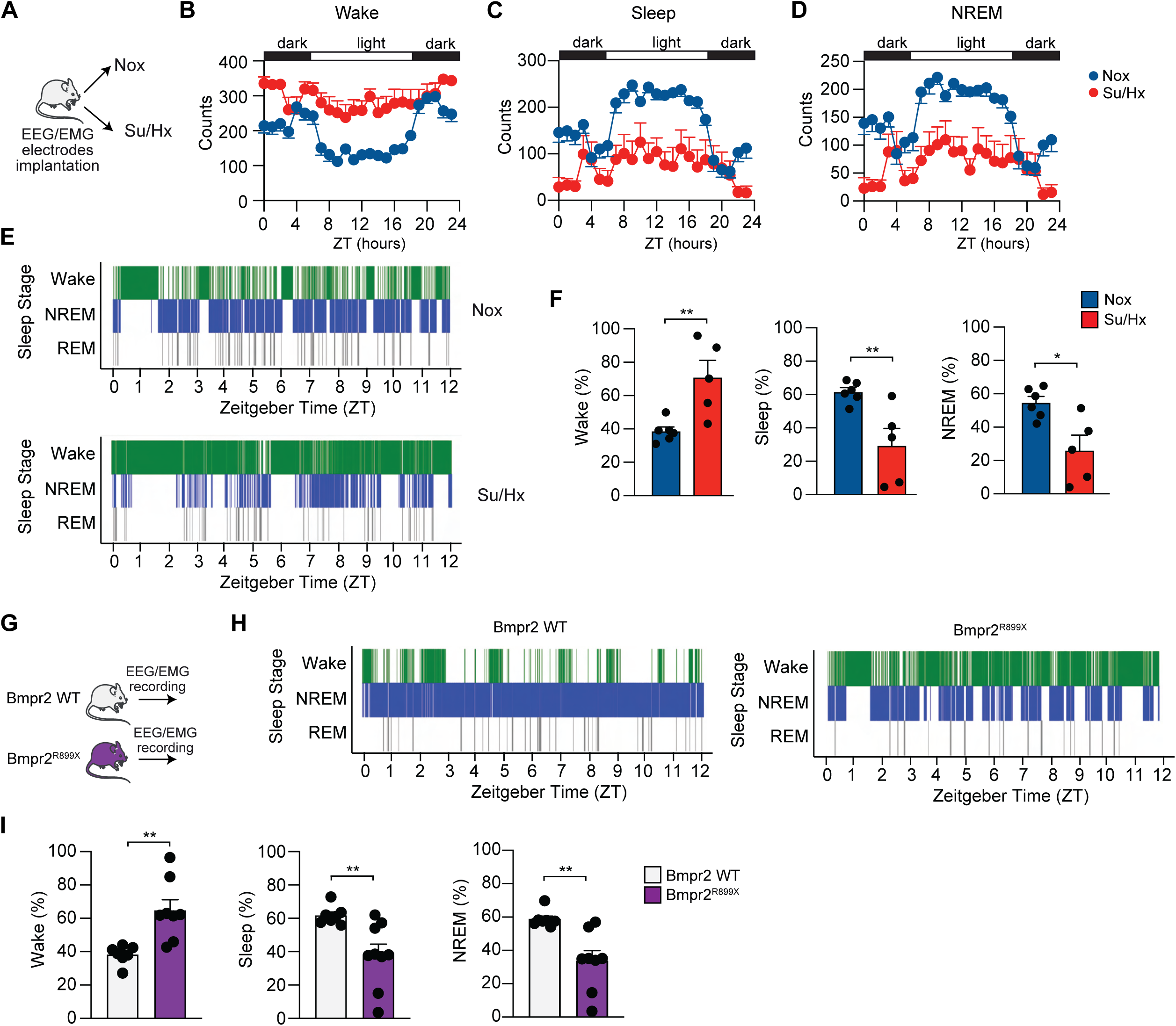
PAH causes poor sleep in mice. (**A**) Study design. Mice were implanted with a headmounted wireless amplifier with three-channel wireless EEG/EMG system and were then exposed to normoxia or Sugen/hypoxia for 4 weeks. (**B-D**) Counts of wake (**B**), sleep (**C**) and Non-Rapid Eye Movement (NREM) (**D**) over an uninterrupted 24 hour period in light/dark cycle. Sleep states were quanti-fied in 10 second epochs. (**E**) Representative hypnograms recorded during the light cycle in mice exposed to normoxia or Sugen/hypoxia conditions. (**F**) Quantification of wake, sleep and NREM states recorded during the light cycle. n =5-6 mice/group. (**G**) Study design. WT and Bmpr2^R899X^ mutant mice were fed a Dox diet during 2 months and were then implanted with a headmounted wireless amplifier with wireless EEG/EMG system. (**H**) Representative hypnograms recorded during the light cycle. (**I**) Quantification of wake, sleep and NREM states recorded during the light cycle. n = 7-9 mice/group.Data were analyzed by two-tailed t-test. Nox: normoxia, Su/Hx: Sugen/Hypoxia.

To verify that poor sleep in the Su/Hx mice is not solely caused by the used PAH model, we assessed sleep quality in smooth muscle-specific doxycycline-inducible BMPR2 mutant mice (SM22-rtTAxTet-BMPR2^R899X^, Figure 7G). These mice spontaneously develop PAH due to a BMPR2 mutation in smooth muscle cells^31^. EEG/EMG analysis of the hypnograms showed that BMPR2 mutant mice have more wake episodes and fewer NREM episodes in comparison to their WT littermates (Figure 7H). Similar to the results obtained using the Su/Hx model, the percentage of time spent awake was significantly higher in BMPR2 mutant mice, with less time in sleep and NREM (Figure 7I). These results demonstrate that PAH caused by Bmpr2 mutation induces poor sleep in mice.

### Improving sleep and reducing inflammation attenuate PAH in mice

Given the decreased sleep quality in PAH and its consequences on lung inflammation, we hypothesized that in such pathological setting improving sleep and dampening inflammation might reduce pulmonary vascular remodeling, ameliorate cardiac dysfunction, and mitigate PAH severity. To test this hypothesis, we improved sleep quality (via a daily injection of melatonin) with or without macrophages depletion (via a weekly injection of Clodronate) in Su/Hx mice during 3 weeks (Figure 8A). Melatonin treatment alone modestly reversed cardiac and pulmonary vascular remodeling (Figure 8B-E). The combination of melatonin and clodronate further markedly reduced RV pressure (Figure 8B) and cardiomyocyte hypertrophy (Figure 8C-D), and significantly attenuated medial thickening of the pulmonary arteries (Figure 8E). Su/Hx mice showed a significant increase in CD68 expression, which was partially reduced by melatonin treatment alone and dramatically decreased by the combination of melatonin with clodronate (Figure 8F). Finally, immunostaining for ki67 revealed decreased PASMC proliferation in the lungs of mice treated with the combination of melatonin and clodronate (Figure 8G). Together, these findings indicate that restoring sleep and inhibiting lung inflammation may prove to be therapeutically effective for treating PAH.

**Figure 8:**
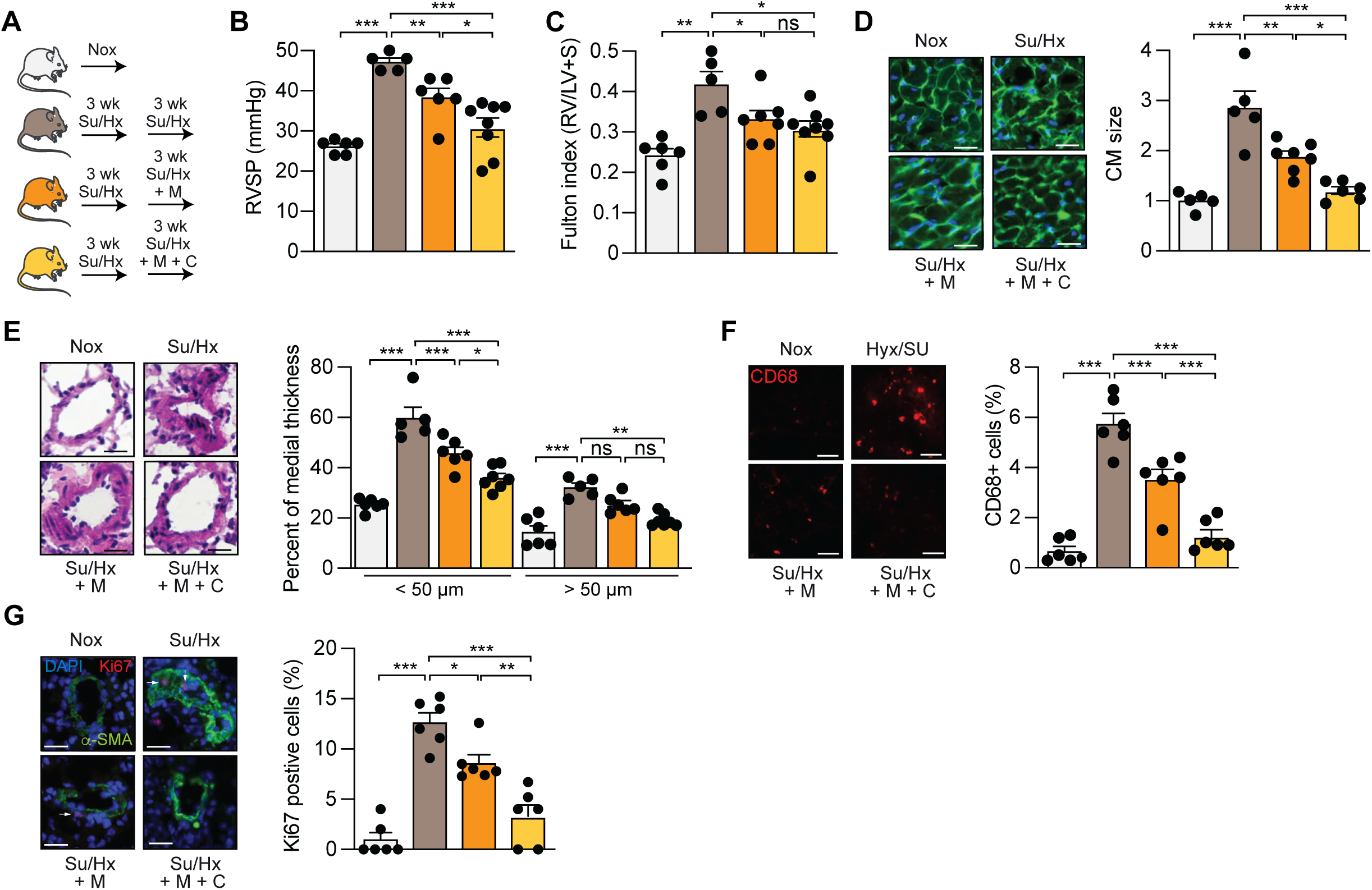
Improving sleep and reducing inflammation attenuate PAH. (**A**) Design of the study. (**B**) RVSP of the indicated groups. (**C**) Fulton index of the indicated groups. (**D**) (Left) Representative WGA-stained RV sections. Scale bar: 50 μm. (Right) Quantitative analysis. (**E**) (Left) Representative H&E-stained pulmonary arteries sections from the indicated groups. Scale bar: 50 μm. (Right) Percentage of arteries medial thickness. (**F**) (Left) Representative lung sections stained with CD68 (red). Scale bar: 50 μm. (Right) Quantitative analysis. (**G**) (Left) Representative lung sections stained with Dapi (blue), α-SMA (green) and Ki-67 (red). Scale bar: 50 μm. (Right) Quantitative analysis. n= 5-8 mice/group. Data were analyzed by one-way ANOVA. Su/Hx: Sugen/Hypoxia; M: melatonin; C: clodronate.

## Discussion

In this study, we identified poor sleep as a novel and modifiable driver of PAH. We used different mouse models of PAH (Su/Hx, Egnl1 KO, IL-6 transgenic, and BMPR2 mutant mice) and two models of sleep disturbances (sleep fragmentation and chronic jet lag) to demonstrate that poor sleep worsens hemodynamic parameters, promotes pulmonary vascular remodeling, and increases right ventricular dysfunction. Importantly, this exacerbation is not caused by physical or psychological stress from the used models. Mechanistically, poor sleep increases the infiltration of macrophages in the lung, disrupts the molecular clock in PASMCs, leading to dysfunctional cell cycle progression and PASMC proliferation. Conversely, inhibiting the major clock driver (Bmal1) or restoring sleep and inhibiting inflammation attenuate PAH. Overall, our findings establish a bidirectional relationship between sleep and PAH, highlighting clock genes, sleep, and immune targets for PAH treatment.

Our findings support the clinical observations reporting prevalence of poor sleep quality and increased sleep disturbance in patients with PAH; characterized by insomnia, sleep fragmentation, and non-restorative sleep^6–10^. These studies consistently reported poor sleep quality in 50% to 73% of patients with PAH and severe insomnia in one-fourth of the PAH patients. While these studies used standardized questionnaires (Pittsburgh Sleep Quality Index, Epworth Sleepiness Scale and Insomnia Severity Index) or wearable sleep trackers, we used an unbiased and accurate method to determine the effects of PAH on sleep quality. EEG/EMG allows the accurate identification of sleep stages and is the gold standard for clinical sleep studies. Using EEG/EMG telemetry, we found that mice with PAH spend most of their time awake during the rest period when compared to healthy mice, indicating that PAH causes poor sleep in mice.

Using multiple experimental models, this study provides direct evidence that poor sleep is not just a comorbidity in PAH but rather a significant factor that contributes to disease severity and progression. In different mouse models of PAH, poor sleep significantly increased right ventricular dysfunction and exacerbated pulmonary vascular remodeling. To exclude the possibility that the observed effects of SF were secondary to stress induced by the SF apparatus rather than sleep fragmentation, we conducted control experiments in which the sweep bar was activated only during the dark cycle (when mice are active and awake). Activation of the SF apparatus when the mice were active did not worsen PAH, indicating that poor sleep (and not mechanical disturbance or stress) underlies the exacerbation of PAH. Additionally, another model of sleep disturbances (i.e., chronic jet lag) confirmed our observations and reinforced the emerging concept that circadian misalignment is a pathogenic driver in cardiovascular diseases^38–42^.

A compelling finding from our study is that poor sleep exacerbates PAH by triggering lung inflammation, specifically macrophage recruitment to the perivascular space. Transcriptomic analyses from the lungs of mice with PAH and exposed to SF revealed significant enrichment of cytokines signaling pathways and leukocyte chemotaxis. Immunofluorescence analysis confirmed increased CD68+ macrophage accumulation in mice with poor sleep. While the role of macrophages has been previously reported in vascular remodeling and PAH^43,44^, our study uniquely links pulmonary macrophage accumulation to poor sleep. In addition, our study demonstrates that inflammatory cytokines upregulate Bmal1 expression in PASMCs. However, it remains unclear how cytokines increase clock genes expression in PASMCs. Interestingly, the absolute level of Bmal1 was increased in PAH-diseased PASMCs, yet its temporal oscillatory dynamics was flattened, and almost lost. This pathological clock uncoupling is similar to that observed in inflammatory diseases and cancers^45–47^. We demonstrate that Bmal1 and Clock control PASMCs functions, as their inhibition decreased PASMCs migration and proliferation while inducing cell cycle checkpoint genes. This is in accordance with previous studies showing that clock genes regulate the cell cycle^48,49^. Consistently, smooth muscle cell-specific deletion of Bmal1 protected mice from Su/Hx-induced PAH, highlighting the cell-specific role of Bmal1 in driving pulmonary vascular remodeling and PAH.

Our findings suggest that PAH itself perturbs sleep-wake regulation and poor sleep exacerbates PAH. Our results demonstrate the presence of a vicious circle in which PAH induces poor sleep, which in turn exacerbates lung inflammation and worsens pulmonary vascular remodeling and PAH. The mechanisms driving poor sleep in PAH remain unclear, although neuroinflammation may play a role. In addition, how poor sleep triggers lung inflammation remains unknown, and future studies should explore the molecular mechanisms of sleep disturbances-induced pulmonary vascular remodeling and PAH. Nevertheless, despite these uncertainties, our findings underscore that circadian clock genes play a crucial role in pulmonary vascular remodeling, at least in part, through their effects on PASMC proliferation and emphasize the potential of inhibiting Bmal1 as a therapeutic target to attenuate PAH.

We explored the therapeutic implications of these findings by administering melatonin to promote sleep and clodronate to deplete macrophages. This therapeutic intervention significantly attenuated PAH by reducing lung inflammation, PASMC proliferation, pulmonary vascular remodeling, and cardiac dysfunction. Melatonin has been historically used to promote sleep, synchronize the circadian rhythm, and as an antioxidant. Lately, melatonin has shown promise in cardiovascular protection^50–52^, in preclinical models of PH^53–55^ and in improving the quality of life of patients with PAH^56^. Our findings provide further rationale for the consideration of using melatonin in combination with an anti-inflammatory drug for the treatment of PAH, particularly for patients with poor sleep.

Although we clearly demonstrated the impact of sleep disturbances on PAH, our study has some limitations. First, we did not perform survival studies to determine whether restoring sleep and reducing inflammation decrease mortality in PAH-diseased animals. Second, we demonstrated the key role of macrophages; however, the contributions of other immune cells were not examined in this study. Third, we did not test the effects of PAH drugs on sleep quality in PAH-diseased animals. Lastly, no animal model can fully replicate the complex causes of PAH in humans, such as autoimmune, thrombotic, or epigenetic factors; therefore, despite using four different animal models of PAH, we did not test these complex mechanisms in our in vivo studies.

### Conclusions

Our studies have identified poor sleep as a novel and modifiable driver of PAH. Poor sleep is not just a comorbidity in PAH but rather a significant factor that contributes to disease severity and progression. We revealed the existence of a vicious circle in which PAH induces poor sleep which in turn worsens pulmonary vascular remodeling and PAH. Improving sleep and inhibiting inflammation has shown promising therapeutic effects in a PAH mouse model, laying the groundwork for further exploration of its clinical applications.

## Acknowledgments

The authors gratefully acknowledge the Histology resources and services provided by the FBRI Histology Core Facility at Virginia Tech Carilion. The authors thank the patients who provided lung samples; the investigators and personnel of the Pulmonary Hypertension Breakthrough Initiative.

## SUPPLEMENTAL MATERIAL

### EXPANDED METHODS

#### In vivo interventions

##### Sugen/Hypoxia-induced PH model

The Sugen/Hypoxia model was induced by exposing mice to 4 weeks of hypoxia (10% O₂) in a ventilated hypoxia chamber with a weekly subcutaneous injection of Sugen 5416 (Sugen, 20 mg/kg; MedChemExpress) for 3 weeks. The normoxic control groups were placed in the chamber and maintained under normoxic conditions (20% O₂).

##### Sleep fragmentation

Mice were placed in a sleep fragmentation (SF) chamber (Lafayette Instrument Company). The chamber fragments mice sleep using tactile stimulation from the motorized moving bar that sweeps the floor of the cage every 2 minutes during the light cycle (when mice are sleeping). The sweep bar automatically shuts off during the dark cycle (when mice are active). The control mice were placed in SF chambers with stationary sweep bars. For experiments involving the Sugen/Hypoxia-induced PAH model, mice (8-10 weeks old) were subjected to 16 weeks of SF in the SF chambers and were then exposed to Sugen/Hypoxia. To induce SF in Egln1 and IL-6 Tg+ models, mice (8-10-week-old) were placed in SF chambers for 16 weeks. At the end of the experiment, cardiac hemodynamic measurements were performed under the same circadian time. Tissues were harvested for morphometric, molecular, and histological analyses.

##### Chronic jet lag

To induce chronic jet lag (CJL), mice (8-10-week-old) were placed in circadian cabinets and exposed to a schedule of 6-hour advances of the light cycle every 2 days for 16 weeks. Control mice were exposed to a normal 12:12 light/dark cycle. The circadian cabinets are fully ventilated and equipped with temperature, humidity, and light level sensors. At the end of the experiment, cardiac hemodynamic measurements were performed under the same circadian time. Tissues were harvested for morphometric, molecular, and histological/immunohistochemical analyses.

##### Melatonin and clodronate treatment

Mice (8-10-week-old) were exposed to Sugen/Hypoxia and then received once weekly intraperitoneal injections of clodronate (10mg per injection, Sapphire North America, Cp-Suv-P-005-005) and daily intraperitoneal injections of melatonin (10mg/kg/day, Sigma, M5250-1g) for a total of 3 weeks while still under hypoxia. The melatonin injection was given 1 hour after LIGHT ON (8 AM) to enhance onset of sleep. The control mice were exposed to Sugen/Hypoxia or normoxia for a total of 6 weeks while receiving a sham. At the end of the experiment, the cardiac and hemodynamic parameters were assessed, and tissues were harvested (under the same circadian time) for immunohistological and molecular analyses.

##### Cardiac hemodynamic measurements

Measurement of right ventricular systolic pressure (RVSP) was performed using the ADVantage Pressure-Volume (PV) Loop System. The system includes a pressure catheter for measuring pressure, which provides real-time feedback about catheter position to ensure quality measurements from every procedure. To perform this procedure, mice were anesthetized with isoflurane and placed on a heated platform for temperature regulation. Then, tracheostomy was performed to expose the trachea and then connected to a small-animal ventilator. A 1.4F pressure-volume catheter was advanced through an opening in the upper abdomen and inserted into the right ventricle. RVSP was measured using the PV loop system to continuously record the pressure waveform during the cardiac cycle. The RVSP was identified as the peak systolic pressure on the waveform. The animals were monitored throughout the procedure for anesthesia depth and recovery.

##### Electroencephalography (EEG) and electromyography (EMG)

To record sleep/wake quality, mice were implanted with wireless electroencephalography (EEG) and Electromyography (EMG) recording electrodes (Pinnacle Technologies). Surgical procedures were performed under anesthesia (2% Isoflurane) using standard aseptic surgical technique with a stereotactic apparatus. The skin was incised (a 2-3 cm incision) along the dorsal midline of the eyes to midway between the scapulae posteriorly, exposing the skull. The connective tissue was removed, and the skull was cleaned with betadine and 100% ethanol. A microdrill was used to make four holes in the skull above the frontal and parietal cortices to place EEG electrodes on the mice. A headmount containing a plastic 6-pin connector attached to four EEG electrodes and ends of two stainless steel Teflon-coated wires serving as EMG electrodes was placed on the skull. Four stainless steel screws were used to anchor the headmounts such that the ends of the screws contact dura of the cortices and serve as EEG recording electrodes. Next, the two Teflon-coated wire ends were inserted into the nuchal musculature on the dorsal of the neck, serving as the EMG electrodes. Sutures were used to close the skin incision and the headmount (and screws) were secured with Palacos dental cement. Mice received postoperative analgesia (ketoprofen) and care and were supplemented with warmth until they fully recovered from the anesthesia. All mice were allowed to recover undisturbed for at least 7 days postoperatively. Mice were placed in SF or control chambers, and EEG/EMG data were collected continuously using the Pinnacle Acquisition software before being analyzed. Data were scored in 10-second epochs per hour as wake, sleep, REM, or NREM counts using IntelliSleepScorer^1^, an automated machine learning assisted scoring program (supplemented with visual confirmation).

##### Morphometric analysis and tissue harvest

After completing the cardiac hemodynamic measurements, mice were perfused with phosphate-buffered saline (PBS) to remove blood. Tissues were immediately harvested under the same circadian time for morphometric and histological/immunohistochemical analyses. The right ventricle (RV) of the heart was separated from the left ventricle plus the septum (LV+S). The ratio of the RV weight and that of the LV+S was calculated to evaluate the Fulton Index, an indicator of right ventricular hypertrophy. Right ventricles and lungs were collected for qPCR, western blot, and immunostaining analyses. Lungs used for immunohistochemical staining were inflated with and embedded in OCT.

##### Hematoxylin & Eosin staining

Lung tissue sections (8 µm) were obtained from OCT-embedded frozen samples and mounted onto glass slides. The slides were then processed for H&E staining to assess the wall thickness of pulmonary arteries. Briefly, sections were air-dried and fixed in 4% paraformaldehyde (PFA) for 15 minutes. After fixation, the sections were stained with Hematoxylin for 10 min, followed by differentiation in 1% acid alcohol and counterstaining with Eosin for 2 min. The slides were then dehydrated through a series of alcohols, cleared in xylene, and mounted with a coverslip. Images were captured using a light microscope, and arterial wall thickness was measured using ImageJ software. Percentage Medial Thickness (% MT) was calculated using the formula;

External Area – Internal area/External area X 100.

To quantify the number of occluded pulmonary arteries (PA) in the lungs, small arteries (<50µm) were counted in the lung sections and assessed for occlusions and scored as: no evidence of occlusion (open), partially occluded (50% closed), or fully occluded (completely closed).

##### Wheat germ agglutinin staining

Right ventricle (RV) tissue sections (8 µm) were obtained from OCT-embedded frozen samples and mounted on glass slides. To preserve tissue morphology, the sections were fixed in 4% paraformaldehyde for 15 minutes at room temperature. After fixation, the sections were washed in PBS to remove excess fixative. WGA conjugated to Alexa Fluor-488 (Thermo Fisher Scientific) was applied to the tissue sections at 1 mg/mL concentration and incubated in a humidified chamber at 37°C for 90 minutes. WGA binds to sialic acid and N-acetylglucosamine residues on cell membranes, specifically highlighting the membrane architecture of cardiomyocytes. After incubation, the sections were washed in PBS to remove unbound WGA. To counterstain the nuclei, the sections were incubated with DAPI (10 µg/mL; Invitrogen) for 5 minutes at room temperature. Finally, the sections were mounted with a coverslip using an anti-fade mounting medium (Vector Laboratories). Images were captured using a fluorescence microscope (Keyence), and the RV tissue morphology was analyzed for hypertrophy with ImageJ.

##### Immunofluorescence

Lung tissue sections (8 µm) were obtained from OCT-embedded frozen samples and mounted onto glass slides. Sections were fixed in 4% paraformaldehyde for 10 min at room temperature (RT), followed by washing with 1X PBS. After blocking with normal goat serum (10%) for 1 hour at RT to prevent non-specific binding, the sections were incubated overnight at 4°C with primary antibodies, BMAL1, CD68, α-SMA, IL-6, Ki-67, and CCR2 (see Table S5). After primary antibody incubation, sections were washed in 1X PBS and incubated with species-specific Alexa Fluor-conjugated secondary antibodies (Invitrogen) for 1 hour at RT in the dark. Nuclei were counterstained with DAPI (10 µg/mL, Invitrogen) for 10 min at RT. After the final PBS washes, sections were mounted with an anti-fade mounting medium and coverslipped. Fluorescent images were captured using a fluorescence microscope (Keyence). K-i67 staining was used to quantify proliferating cells, IL-6 was used to assess cytokine, and CD68 and CCR2 for macrophage expression in the lung. α-SMA was used as a marker of smooth muscle cells. Image analysis was performed using ImageJ to quantify respective protein expressions. To calculate the percentage of CD68+ and CCR2+ positive cells in the lung sections, positive cells were counted using thresholding and particle analysis in at least five representative images per sample. The counts were normalized to the number of DAP-stained nuclei. To calculate the percentage of Ki-67 positive cells, Ki-67+ nuclei within α-SMA-stained regions in the pulmonary arteries were identified and counted using thresholding and particle analysis across five representative images per sample. The counts were normalized to DAPI+ nuclei within the same α-SMA+ regions. The α-SMA+ regions were defined by region of interest (ROI) drawing. The IL-6 intensity per cell was calculated by measuring the integrated density in at least five representative images per sample, and the signal was normalized to the number of DAP-stained nuclei. The BMAL1 intensity per unit area was calculated by measuring the integrated density of BMAL1 signal within the ROI in at least five representative images per sample, and normalized to the area of the artery.

##### Cell culture

Human pulmonary arterial smooth muscle cells (PASMCs) and endothelial cells (PAECs) were obtained from Lonza, as well as from the Pulmonary Hypertension Biobank Initiative (PHBI). Cells from female and male patients were used for the study, and experiments were performed at least 5 times with a minimum of 3 replicates. PASMCs and PAECs were cultured with VascuLife Basal medium (SMC; lot # 11829) and (VEGF-Mv; lot # 12310), respectively, supplemented with 5% fetal bovine serum (5% FBS) and 1% Ampicillin and Gentamicin, in a humidified incubator at 37°C with 5% CO2 in air. All experiments were performed using primary cells between passages two and eight.

##### Synchronization and circadian rhythmicity in cultured PASMCs and PAECs

PASMCS and PAECs were seeded in 6-well plates and allowed to reach 70% confluence before being synchronized by treating with 200 nM Dexamethasone (dissolved in 95% ethanol with 0.1% FBS) or ethanol containing 0.1% FBS as a control for 1 hour. After treatment, the media containing dexamethasone was removed, and the cells were washed with 1x PBS before samples were collected every 4 hours over 24 hours to assess gene and protein expressions of clock genes. These data were later analysed with Cosinor to determine circadian oscillation. Zeitgeber time (ZT0) is defined as the time 0 hours after dexamethasone treatment. To evaluate the effects of inflammatory cytokines and growth factors on clock gene expressions, PAMSCs were synchronized with Dexamethasone and treated with TNF-α (10 ng/ml; F peprotech), IL-1β (10 ng/ml), TGF-β (5 ng/ml), PDGF (10 ng/ml), or ET-1 (1nM) for 48 hours, and samples were collected at ZT12 post synchronization.

##### siRNA transfection

Small interfering RNAs (siRNAs) specifically designed to target Bmal1 (siBmal1), Clock (siClock), and scrambled siRNA (siCtrl) were purchased from Qiagen. Cells were seeded onto 6-well plates, and 5 nM of siRNA was transfected using Lipofectamine RNAi-MAX (Invitrogen). The transfection medium was replaced 6 hours later with fresh VascuLife Basal medium, and cells were harvested after 48 hours for analysis or further treatment. The efficiency of the siRNAs was first confirmed at both the mRNA and protein levels. Additionally, the effects of siBmal1 and siClock on clock genes and cell cycle genes were analysed by qPCR, and relative expressions were compared with those of siCtrl.

##### Proliferation assay

For proliferation assay, PASMCs from healthy donors were seeded in 96-well plates at a concentration of 4000 cells/ well and incubated overnight at 37°C with 5% CO2. One day later, cells were incubated with the different treatments in 0.1% FBS or 5% FBS medium for 48 hours. BrdU labelling solution was then added to each well and incubated for an additional 16 hours, and the proliferation assay subsequently performed following the manufacturer’s instructions. To assess the effect of Bmal1 and Clock knockdown on PASMC proliferation, 2000 cells/well were seeded in 96-well plates overnight, and cells were transfected with siBmal1 and siClock or siCtrl. Forty-eight hours after transfection, a BrdU labelling solution was added to each well, and a proliferation assay was performed according to the manufacturer’s instructions.

##### Wound healing assay

PASMCs from healthy donors were seeded in a 12-well plate at a high density (80%-90% confluence). One day later, the cells were transfected with siRNAs designed against Bmal1, Clock, or a scrambled sequence. Two days later, scratching was performed and examined under a microscope. After removing any detached cells by washing with PBS, 1 mL of 0.1% FBS or 5% fresh culture medium was added to the wells. Afterwards, plates were incubated, and wells were then photographed at different time points (0 h and 48 h). ImageJ software was used to measure the width of the scratch at different time points. To calculate the percentage of wound area (to denote the percentage of cells that repopulate the wound), the following formula was used:

Wound area (%) = (Initial Wound Width-Final Wound Width/Initial Wound Width) ×100

##### RNA extraction

Total RNA was extracted from cells or ground tissues using the TRIzol reagent (Zymo Research), following the manufacturer’s protocol. Briefly, samples were homogenized in TRIzol, and after phase separation, the aqueous phase containing RNA was carefully collected. RNA was then precipitated using isopropanol, washed with 75% ethanol, and resuspended in RNase-free water. To remove any residual DNA, RNA samples were treated with DNase I (Thermo Scientific) following the manufacturer’s instructions. RNA quantity and quality were assessed using a NanoDrop spectrophotometer (Thermo Scientific), and samples were stored at −80°C until further analysis.

##### Reverse transcription

For reverse transcription, 500 ng of total RNA was used as the template for cDNA synthesis using the High-Capacity cDNA Reverse Transcription Kit (Applied Biosystems), following the manufacturer’s protocol. The reaction was performed at 25°C for 10 min, 37°C for 120 min, and 85°C for 5 min to inactivate the reverse transcriptase. The resulting cDNA was stored at −20°C for downstream quantitative PCR (qPCR) analysis.

##### Real-time quantitative PCR

Quantitative PCR (qPCR) was performed using SYBR Green Master Mix (Applied Biosystems) and gene-specific primers for target genes. The reaction conditions included an initial denaturation at 95°C for 10 min, followed by 40 cycles of denaturation at 95°C for 15 sec, annealing at 60°C for 30 sec, and extension at 72°C for 30 sec. Melting curve analysis was performed after amplification to confirm primer specificity. Gene expression levels were normalized to housekeeping genes (GAPDH or β-actin), and the relative expression of target genes (see table S4 for primers) was calculated using the ΔΔCt method, where the Ct value of each gene in the experimental group was compared to the control group to determine fold changes in expression.

##### RNA-seq

For RNA sequencing (RNA-Seq), total RNA was prepared as described above, and quality was further assessed using the Agilent Bioanalyzer (Agilent Technologies) to ensure RNA integrity (RIN value >7). High-quality RNA (≥1 µg) was sent to Novogene for library preparation and sequencing. Briefly, RNA samples were enriched for mRNA using poly(A) selection, and library construction was performed using the Illumina TruSeq Stranded mRNA Library Prep Kit. The libraries were then sequenced on the Illumina NovaSeq platform, generating high-depth, paired-end reads. The raw sequencing data were processed and analyzed by Novogene to obtain gene expression profiles, which were used for downstream bioinformatics analysis.

##### Statistical Analysis

All data are presented as mean ± standard error of the mean (SEM). Statistical analyses were performed using GraphPad Prism. Differences between two groups were analyzed using an unpaired two-tailed Student’s t-test, while comparisons among multiple groups were conducted using one-way or two-way ANOVA, followed by Tukey’s post hoc test, as appropriate. The normality of data distribution was assessed using the Shapiro-Wilk test. A p-value < 0.05 was considered statistically significant.

## SUPPLEMENTAL TABLES

**Table S1.** Detailed PASMC characteristics used in this study.

|  | **Sex** | **Age** | **Race** | **PAH** |
| --- | --- | --- | --- | --- |
| **Control 1** | Female | 56 | Caucasian | - |
| **Control 2** | Male | 46 | Caucasian | - |
| **Control 3** | Female | 47 | Caucasian | - |
| **Control 4** | Male | 44 | Caucasian | - |
| **Control 5** | Male | 45 | Caucasian | - |
| **Control 6** | Female | 36 | Caucasian | - |
| **Control 7** | Male | 49 | Caucasian | - |
| **PAH 1** | Male | 40 | Caucasian | iPAH |
| **PAH 2** | Female | 33 | Caucasian | iPAH |
| **PAH 3** | Female | 58 | Caucasian | iPAH |
| **PAH 4** | Male | 45 | Caucasian | iPAH |
| **PAH 5** | Female | 47 | Caucasian | iPAH |
| **PAH 6** | Male | 43 | Caucasian | iPAH |
| **PAH 7** | Female | 55 | Caucasian | iPAH |

**Table S2.** Detailed PAEC characteristics used in this study.

|  | **Sex** | **Age** | **Race** | **PAH** |
| --- | --- | --- | --- | --- |
| **Control 1** | Female | 57 | Caucasian | - |
| **Control 2** | Male | 49 | Caucasian | - |
| **Control 3** | Female | 55 | Caucasian | - |
| **Control 4** | Female | 50 | Caucasian | - |
| **Control 5** | Male | 25 | Caucasian | - |
| **Control 6** | Female | 49 | Caucasian | - |
| **PAH 1** | Male | 40 | Caucasian | iPAH |
| **PAH 2** | Female | 32 | Caucasian | iPAH |
| **PAH 3** | Female | 27 | Caucasian | iPAH |
| **PAH 4** | Female | 40 | Caucasian | iPAH |
| **PAH 5** | Male | 51 | Caucasian | iPAH |
| **PAH 6** | Male | 40 | Asian | iPAH |

**Table S3.** Detailed characteristics of the PAH patients.

|  | **Sex** | **Age** | **Race** | **PAH** |
| --- | --- | --- | --- | --- |
| **Control 1** | Female | 28 | Caucasian | - |
| **Control 2** | Male | 33 | Hispanic | - |
| **Control 3** | Male | 44 | Caucasian | - |
| **Control 4** | Male | 54 | Hispanic | - |
| **Control 5** | Male | 26 | Caucasian | - |
| **Control 6** | Female | 43 | Caucasian | - |
| **Control 7** | Female | 34 | Caucasian | - |
| **PAH 1** | Female | 55 | Caucasian | iPAH |
| **PAH 2** | Male | 29 | Asian | iPAH |
| **PAH 3** | Male | 25 | Caucasian | iPAH |
| **PAH 4** | Male | 40 | Caucasian | iPAH |
| **PAH 5** | Male | 26 | Caucasian | iPAH |
| **PAH 6** | Female | 32 | Caucasian | iPAH |
| **PAH 7** | Male | 53 | Caucasian | iPAH |

**Table S4.**
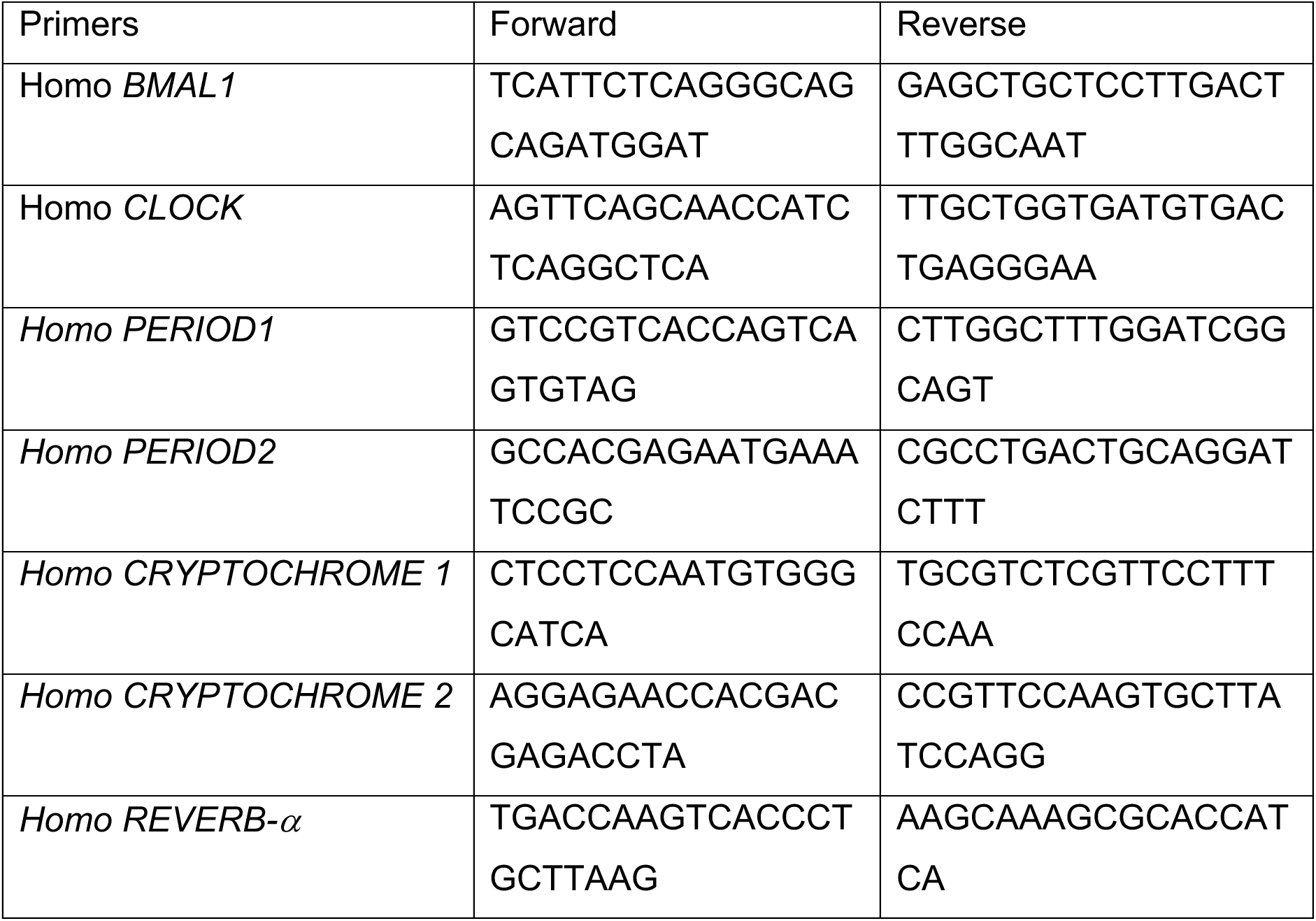
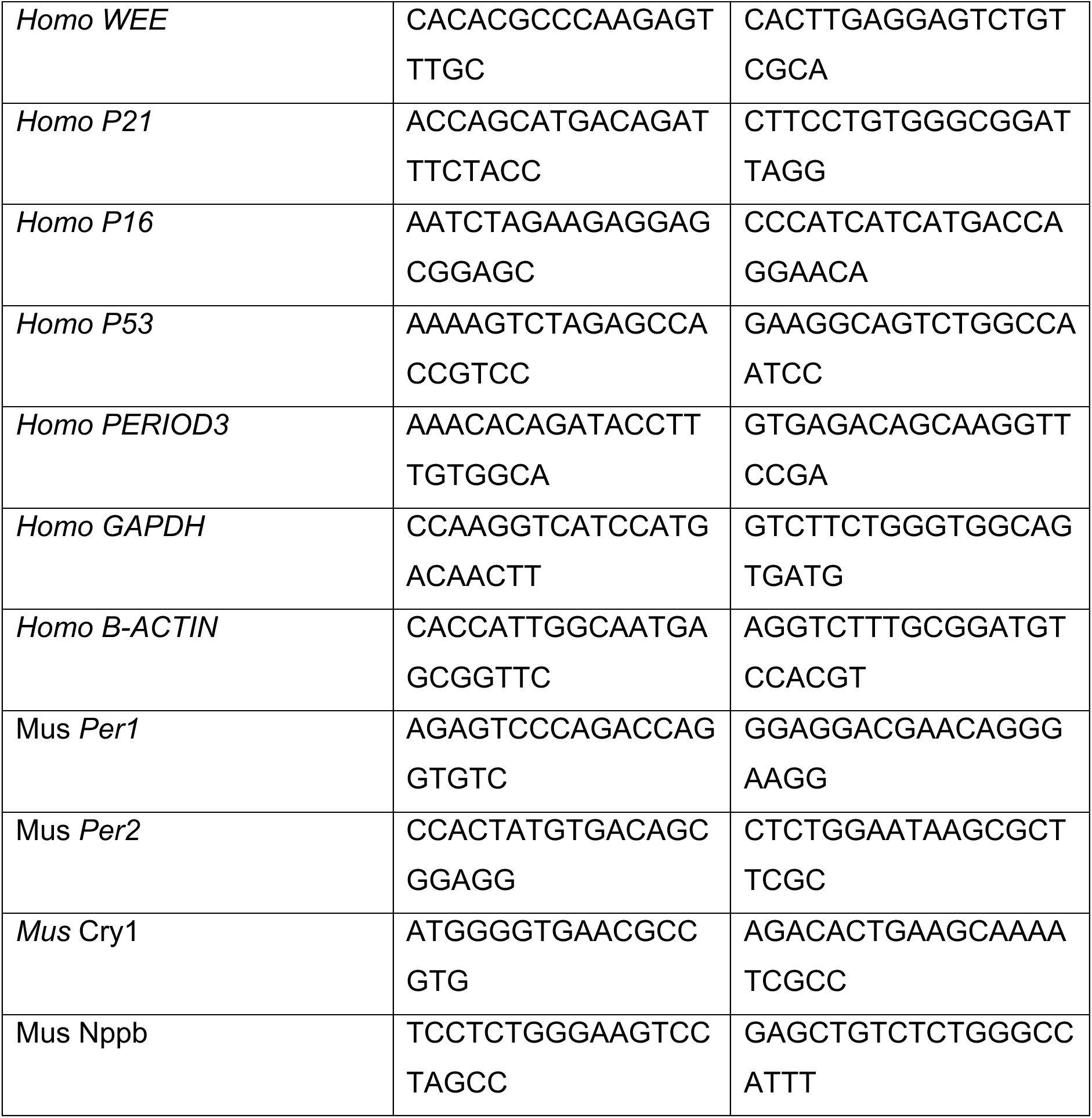
Lists of Primers.

**Table S5.** List of antibodies.

| Antibodies | Vendors | Catalog no. |
| --- | --- | --- |
| ki67 | Abcam | ab16667 |
| CD68 | Invitrogen | 14-0681-82 |
| α-SMA | Cell Signaling | 34105S |
| BMAL1 | Abcam | ab93806 |
| CCR2 | Thermofisher | PA5-23043 |
| IL6 | Invitrogen | P620 |

## Supplemental Figures

**Figure SI:**
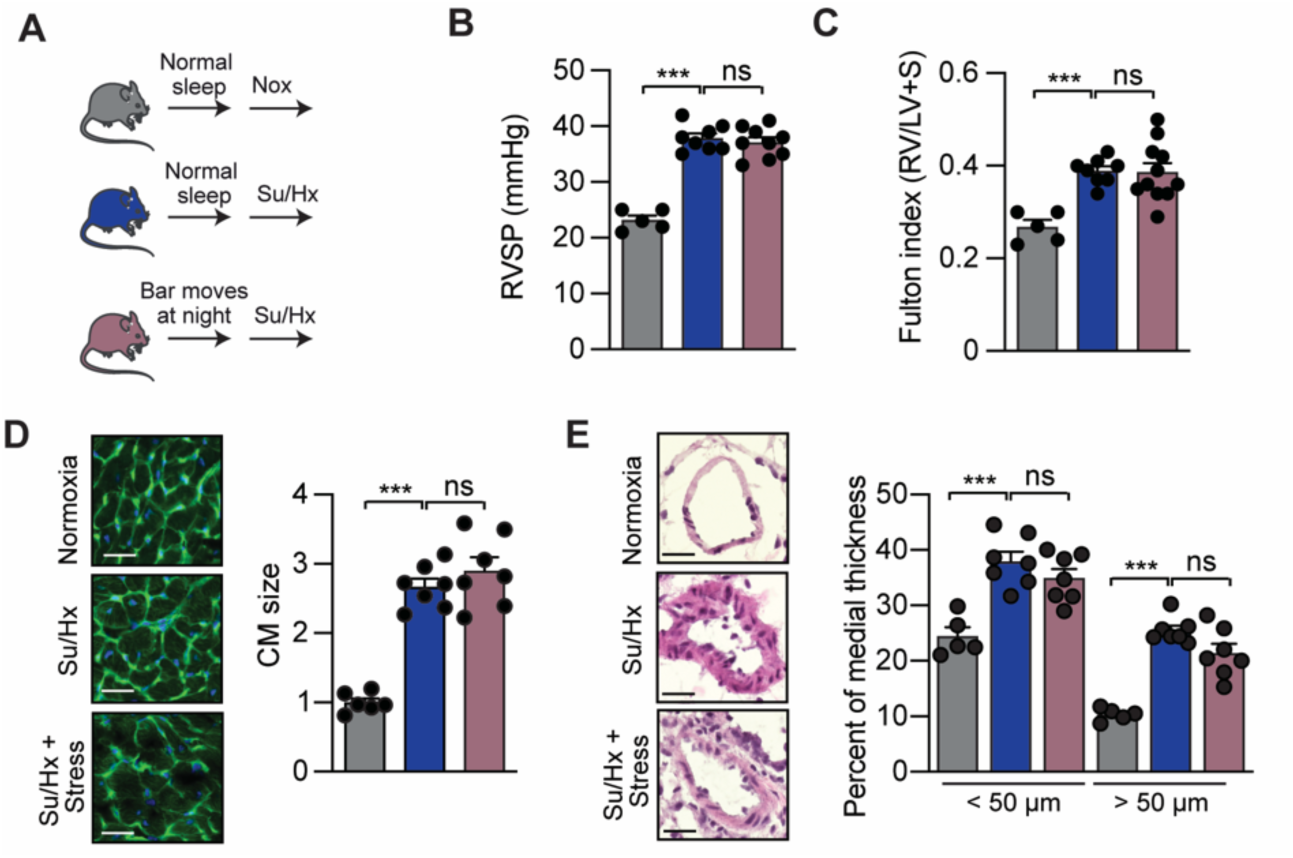
Stress do not exacerbates PAH. **(A)** Experimental design. **(B)** RVSP of the indicated groups, **(C)** Fulton index of the indicat­ed groups. (D) (Left) Representative WGA-stained RV sections. Scale bar: 50 µm. (Right) Quantitative analysis. (E) (Left) Representative H&E-stained pulmonary arteries sections from the indicated groups. Scale bar: 50 µm. (Right) Percentage of arteries medial thickness, n= 5-9 mice/group. Data were analyzed by 1-way ANOVA. *P<0.05, ** P<0.01, ***P<0.001.

**Figure S2:**
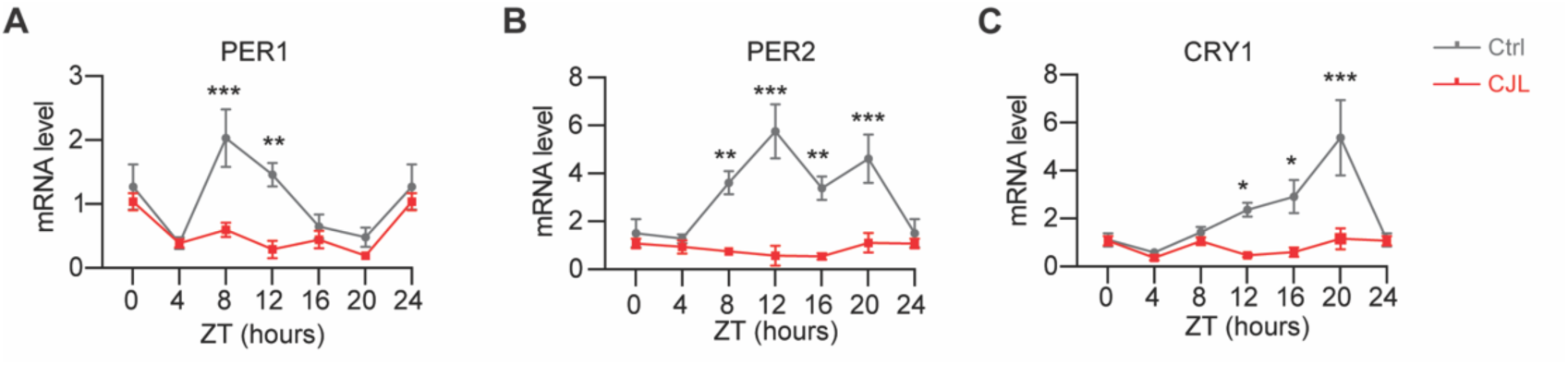
The chronic jet lag model of sleep disturbance. **(A)** Light regime for chronic jet lag treatment. Male wildtype mice were exposed to stable or shiftwork-like light conditions for 64 days. Black bars represent dark phase and hollow bars represent light phase. **(A)** Per1, **(B)** Per 2 and **(C)** Cry1 mRNA levels in the lungs of control or CJL-exposed mice. n = 5 mice/time poinUgroup.

**Figure S3:**
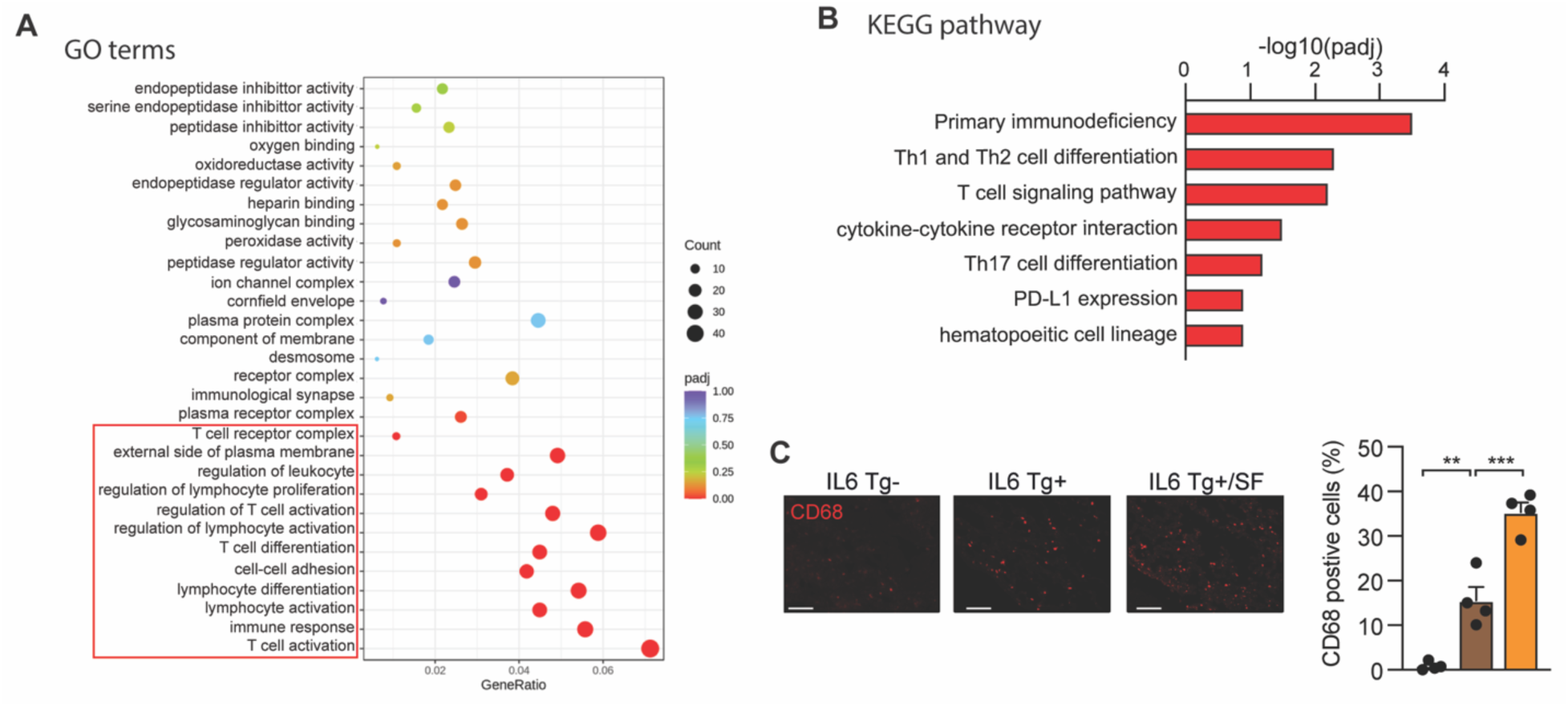
Poor sleep triggers inflammation in IL-6 transgenic mice. **(A)** Scatter plot of top 30 gene ontology (GO) terms enriched by differentially expressed genes (DEGs) in Su/Hx+SF vs Su/Hx lungs (n=4 mice/group). ‘padj’ is the P-value adjusted using the Benjami­ ni-Hochberg procedure. ‘Count’ is the number of genes enriched in a GO term. ‘GeneRatio’ is the percentage of total DEGs in the given GO term. **(B)** KEGG pathway enrichment analysis.The significant pathway for differentially expressed genes in Su/Hx+SF vs Su/Hx lungs. **(C)** CD68+ cells (red) in the respective lungs with quantification. Scale bar: 20 µm. •• P<0.01, •••P<0.001. Data were analyzed by 1-way ANOVA.

**Figure S4:**
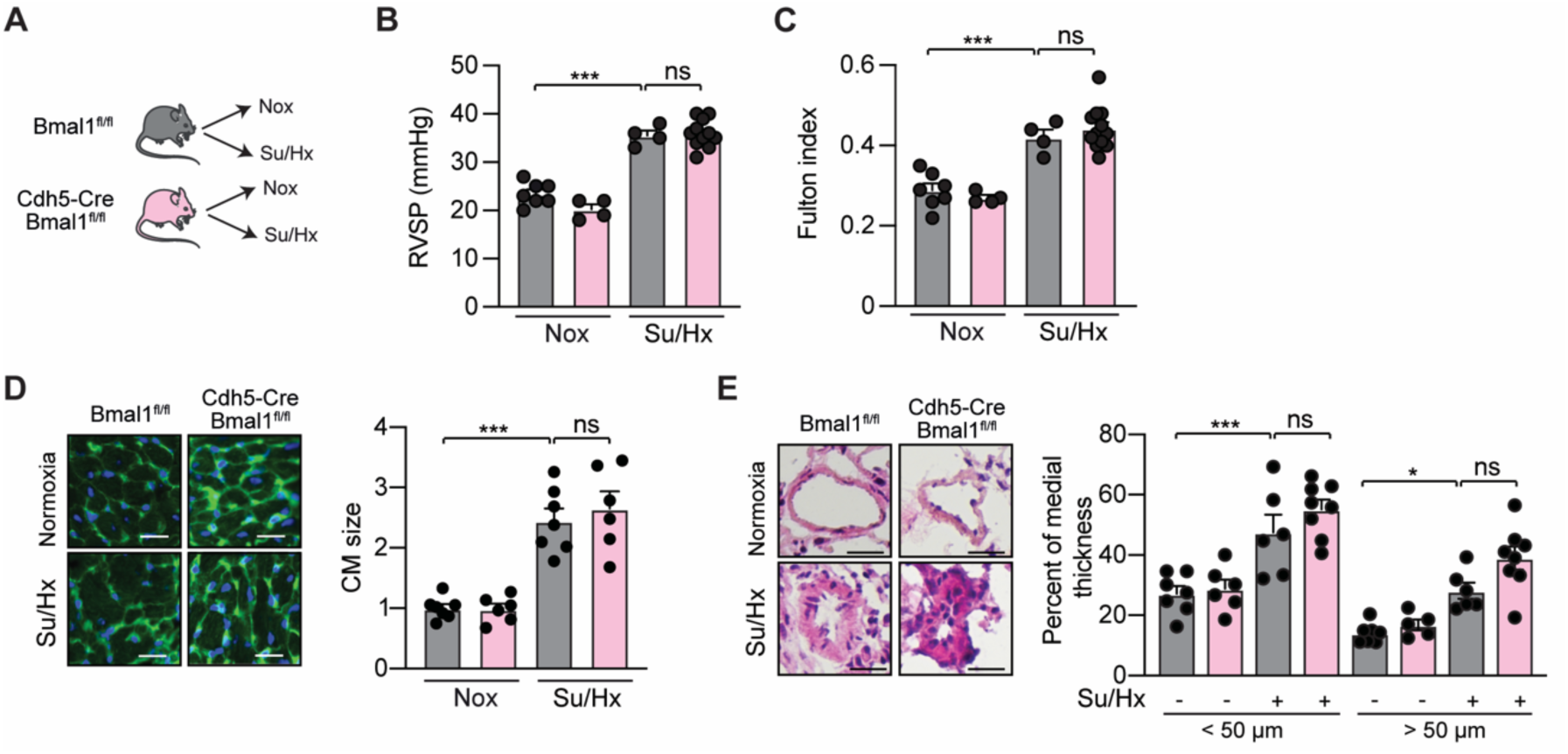
EC-specific Bmal1 deletion do not protects mice against PAH. **(A)** Design of the study. **(B)** RVSP of the indicated groups. **(C)** Fulton index of the indicated groups. **(D)** (Left) Representative WGA-stained RV sections. Scale bar: 50 µm. (Right) Quantitative analysis. **(E)** (Left) Representative H&E-stained pulmonary arteries sections from the indicated groups. Scale bar: 50 µm. (Right) Percentage of arteries medial thickness. n= 6-7 mice/group. Data were analyzed by 2-way ANOVA.

**Figure S5:**
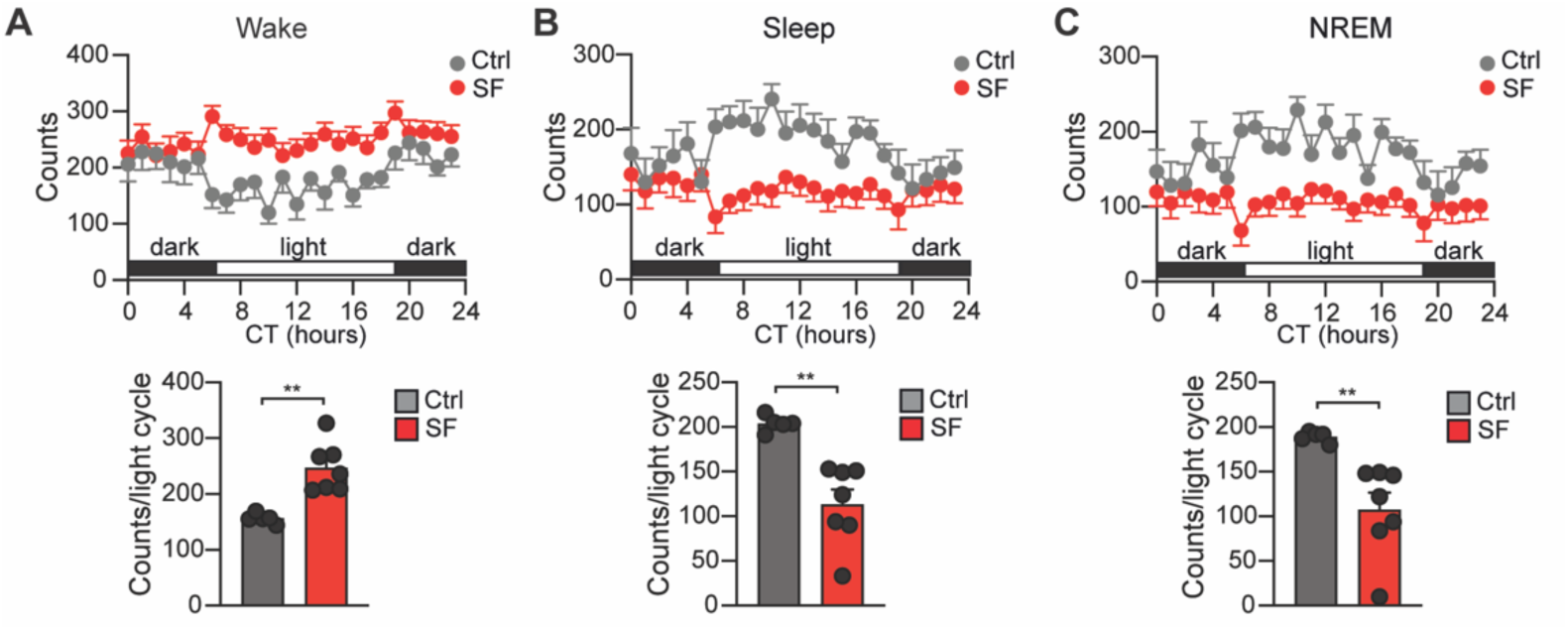
EEG/EMG telemetry to measure sleep disturbances in mice. **(A)** A three-channel wireless EEG/EMG system was used to record wake, sleep and NREM states in control mice and mice under sleep fragmentation. Mice were implanted with a headmounted wireless amplifier that streams data to a computer in real-time. The state of sleep, wake or NREM was determined every 2 minutes in the used mice. **(A)** (Upper panel) Quantification of wake state over an uninterrupted 24 hour period in light/dark cycle. (Lower panel) Total wake counts recorded during the light cycle. **(B)** (Upper panel) Quantification of sleep state in control or SF mice over a 24 h period in light/dark cycle. (Lower panel) Total sleep counts recorded during the light cycle. (C) (Upper panel) Quantification of Non-Rapid Eye Movement (NREM) state in control or SF mice over a 24 h period in light/dark cycle. (Lower panel) Total NREM counts recorded during the light cycle. n = 5-7 mice/group. Data were analyzed by two-tailed t-test.

## References

1. Humbert M, Sitbon O, Chaouat A, Bertocchi M, Habib G, Gressin V, Yaici A, Weitzenblum E, Cordier JF, Chabot F, et al. Survival in patients with idiopathic, familial, and anorexigen-associated pulmonary arterial hypertension in the modern management era. Circulation. 2010;122:156–163. doi: 10.1161/CIRCULATIONAHA.109.911818

2. Hassoun PM. Pulmonary Arterial Hypertension. N Engl J Med. 2021;385:2361-2376. doi: 10.1056/NEJMra2000348

3. Chang KY, Duval S, Badesch DB, Bull TM, Chakinala MM, De Marco T, Frantz RP, Hemnes A, Mathai SC, Rosenzweig EB, et al. Mortality in Pulmonary Arterial Hypertension in the Modern Era: Early Insights From the Pulmonary Hypertension Association Registry. J Am Heart Assoc. 2022;11:e024969. doi: 10.1161/JAHA.121.024969

4. Boucherat O, Bonnet S, Provencher S, Potus F. Anti-remodeling therapies in pulmonary arterial hypertension. Trends Pharmacol Sci. 2025;46:674–691. doi: 10.1016/j.tips.2025.05.004

5. Sitbon O, Boucly A, Weatherald J, Antigny F, Guignabert C, Jevnikar M, Jaïs X, Price LC, Wilkins MR, Zamanian RT, et al. Drugs targeting novel pathways in pulmonary arterial hypertension. Eur Respir J. 2025. doi: 10.1183/13993003.01830-2024

6. Swinnen K, Delcroix M, Belge C, Quarck R, Testelmans D, Buyse B, Helsen F. Impact of insomnia on exercise capacity and quality of life in patients with pulmonary arterial hypertension. European Respiratory Journal. 2017;50. doi: 10.1183/1393003.congress-2017.PA2418

7. Batal O, Khatib OF, Bair N, Aboussouan LS, Minai OA. Sleep quality, depression, and quality of life in patients with pulmonary hypertension. Lung. 2011;189:141–149. doi: 10.1007/s00408-010-9277-9

8. Tiede H, Rorzyczka J, Dumitrascu R, Belly M, Reichenberger F, Ghofrani HA, Seeger W, Heitmann J, Schulz R. Poor sleep quality is associated with exercise limitation in precapillary pulmonary hypertension. BMC Pulm Med. 2015;15:11. doi: 10.1186/s12890-015-0005-3

9. Matura LA, McDonough A, Hanlon AL, Carroll DL, Riegel B. Sleep disturbance, symptoms, psychological distress, and health-related quality of life in pulmonary arterial hypertension. Eur J Cardiovasc Nurs. 2015;14:423–430. doi: 10.1177/1474515114537951

10. Hughes AM, Lindsey A, Annis J, Burke K, Master H, Silverman-Lloyd LG, Garry JD, Blaha MJ, Berman Rosenzweig ES, Frantz RP, et al. Physical Activity, Sleep, and Quality of Life in Pulmonary Arterial Hypertension: Novel Insights From Wearable Devices. Pulm Circ. 2025;15:e70069. doi: 10.1002/pul2.70069

11. Haspel JA, Anafi R, Brown MK, Cermakian N, Depner C, Desplats P, Gelman AE, Haack M, Jelic S, Kim BS, et al. Perfect timing: circadian rhythms, sleep, and immunity - an NIH workshop summary. JCI Insight. 2020;5. doi: 10.1172/jci.insight.131487

12. Rijo-Ferreira F, Takahashi JS. Genomics of circadian rhythms in health and disease. Genome Med. 2019;11:82. doi: 10.1186/s13073-019-0704-0

13. Hunt T, Sassone-Corsi P. Riding tandem: circadian clocks and the cell cycle. Cell. 2007;129:461–464. doi: 10.1016/j.cell.2007.04.015

14. Hou T, Su W, Duncan MJ, Olga VA, Guo Z, Gong MC. Time-restricted feeding protects the blood pressure circadian rhythm in diabetic mice. Proc Natl Acad Sci U S A. 2021;118. doi: 10.1073/pnas.2015873118

15. Buhr ED, Takahashi JS. Molecular components of the Mammalian circadian clock. Handb Exp Pharmacol. 2013:3–27. doi: 10.1007/978-3-642-25950-0_1

16. Astiz M, Heyde I, Oster H. Mechanisms of Communication in the Mammalian Circadian Timing System. Int J Mol Sci. 2019;20. doi: 10.3390/ijms20020343

17. Richards J, Gumz ML. Advances in understanding the peripheral circadian clocks. FASEB J. 2012;26:3602–3613. doi: 10.1096/fj.12-203554

18. Zhang S, Dai M, Wang X, Jiang SH, Hu LP, Zhang XL, Zhang ZG. Signalling entrains the peripheral circadian clock. Cell Signal. 2020;69:109433. doi: 10.1016/j.cellsig.2019.109433

19. Bunger MK, Wilsbacher LD, Moran SM, Clendenin C, Radcliffe LA, Hogenesch JB, Simon MC, Takahashi JS, Bradfield CA. Mop3 is an essential component of the master circadian pacemaker in mammals. Cell. 2000;103:1009–1017. doi: 10.1016/s0092-8674(00)00205-1

20. Menet JS, Pescatore S, Rosbash M. CLOCK:BMAL1 is a pioneer-like transcription factor. Genes Dev. 2014;28:8–13. doi: 10.1101/gad.228536.113

21. Takahashi JS. Transcriptional architecture of the mammalian circadian clock. Nat Rev Genet. 2017;18:164–179. doi: 10.1038/nrg.2016.150

22. Ruan W, Yuan X, Eltzschig HK. Circadian rhythm as a therapeutic target. Nat Rev Drug Discov. 2021;20:287–307. doi: 10.1038/s41573-020-00109-w

23. Logan RW, Zhang C, Murugan S, O’Connell S, Levitt D, Rosenwasser AM, Sarkar DK. Chronic shift-lag alters the circadian clock of NK cells and promotes lung cancer growth in rats. J Immunol. 2012;188:2583–2591. doi: 10.4049/jimmunol.1102715

24. Walker WH, 2nd, Walton JC, DeVries AC, Nelson RJ. Circadian rhythm disruption and mental health. Transl Psychiatry. 2020;10:28. doi: 10.1038/s41398-020-0694-0

25. El Jamal N, Lordan R, Teegarden SL, Grosser T, FitzGerald G. The Circadian Biology of Heart Failure. Circ Res. 2023;132:223–237. doi: 10.1161/CIRCRESAHA.122.321369

26. Delisle BP, Prabhat A, Burgess DE, Ono M, Esser KA, Schroder EA. Circadian Regulation of Cardiac Arrhythmias and Electrophysiology. Circ Res. 2024;134:659–674. doi: 10.1161/CIRCRESAHA.123.323513

27. Fishbein AB, Knutson KL, Zee PC. Circadian disruption and human health. J Clin Invest. 2021;131. doi: 10.1172/jci148286

28. Lim AS, Kowgier M, Yu L, Buchman AS, Bennett DA. Sleep Fragmentation and the Risk of Incident Alzheimer’s Disease and Cognitive Decline in Older Persons. Sleep. 2013;36:1027–1032. doi: 10.5665/sleep.2802

29. Zheng NS, Annis J, Master H, Han L, Gleichauf K, Ching JH, Nasser M, Coleman P, Desine S, Ruderfer DM, et al. Sleep patterns and risk of chronic disease as measured by long-term monitoring with commercial wearable devices in the All of Us Research Program. Nat Med. 2024;30:2648–2656. doi: 10.1038/s41591-024-03155-8

30. Steiner MK, Syrkina OL, Kolliputi N, Mark EJ, Hales CA, Waxman AB. Interleukin-6 overexpression induces pulmonary hypertension. Circ Res. 2009;104:236–244, 228p following 244. doi: 10.1161/CIRCRESAHA.108.182014

31. West J, Harral J, Lane K, Deng Y, Ickes B, Crona D, Albu S, Stewart D, Fagan K. Mice expressing BMPR2R899X transgene in smooth muscle develop pulmonary vascular lesions. Am J Physiol Lung Cell Mol Physiol. 2008;295:L744–755. doi: 10.1152/ajplung.90255.2008

32. Dai Z, Li M, Wharton J, Zhu MM, Zhao YY. Prolyl-4 Hydroxylase 2 (PHD2) Deficiency in Endothelial Cells and Hematopoietic Cells Induces Obliterative Vascular Remodeling and Severe Pulmonary Arterial Hypertension in Mice and Humans Through Hypoxia-Inducible Factor-2α. Circulation. 2016;133:2447–2458. doi: 10.1161/circulationaha.116.021494

33. Deaton RA, Bulut G, Serbulea V, Salamon A, Shankman LS, Nguyen AT, Owens GK. A New Autosomal Myh11-CreER(T2) Smooth Muscle Cell Lineage Tracing and Gene Knockout Mouse Model-Brief Report. Arterioscler Thromb Vasc Biol. 2023;43:203–211. doi: 10.1161/ATVBAHA.122.318160

34. Stacher E, Graham BB, Hunt JM, Gandjeva A, Groshong SD, McLaughlin VV, Jessup M, Grizzle WE, Aldred MA, Cool CD, et al. Modern age pathology of pulmonary arterial hypertension. Am J Respir Crit Care Med. 2012;186:261–272. doi: 10.1164/rccm.201201-0164OC

35. Stearman RS, Bui QM, Speyer G, Handen A, Cornelius AR, Graham BB, Kim S, Mickler EA, Tuder RM, Chan SY, et al. Systems Analysis of the Human Pulmonary Arterial Hypertension Lung Transcriptome. Am J Respir Cell Mol Biol. 2019;60:637–649. doi: 10.1165/rcmb.2018-0368OC

36. Shostak A. Circadian Clock, Cell Division, and Cancer: From Molecules to Organism. Int J Mol Sci. 2017;18. doi: 10.3390/ijms18040873

37. Feillet C, van der Horst GT, Levi F, Rand DA, Delaunay F. Coupling between the Circadian Clock and Cell Cycle Oscillators: Implication for Healthy Cells and Malignant Growth. Front Neurol. 2015;6:96. doi: 10.3389/fneur.2015.00096

38. McAlpine CS, Kiss MG, Rattik S, He S, Vassalli A, Valet C, Anzai A, Chan CT, Mindur JE, Kahles F, et al. Sleep modulates haematopoiesis and protects against atherosclerosis. Nature. 2019;566:383–387. doi: 10.1038/s41586-019-0948-2

39. Ziegler KA, Engelhardt S, Carnevale D, McAlpine CS, Guzik TJ, Dimmeler S, Swirski FK. Neural Mechanisms in Cardiovascular Health and Disease. Circ Res. 2025;136:1233–1261. doi: 10.1161/circresaha.125.325580

40. Huynh P, Hoffmann JD, Gerhardt T, Kiss MG, Zuraikat FM, Cohen O, Wolfram C, Yates AG, Leunig A, Heiser M, et al. Myocardial infarction augments sleep to limit cardiac inflammation and damage. Nature. 2024;635:168–177. doi: 10.1038/s41586-024-08100-w

41. Durgan DJ, Young ME. The cardiomyocyte circadian clock: emerging roles in health and disease. Circ Res. 2010;106:647–658. doi: 10.1161/circresaha.109.209957

42. Chellappa SL, Vujovic N, Williams JS, Scheer F. Impact of Circadian Disruption on Cardiovascular Function and Disease. Trends Endocrinol Metab. 2019;30:767–779. doi: 10.1016/j.tem.2019.07.008

43. Ferrian S, Cao A, McCaffrey EF, Saito T, Greenwald NF, Nicolls MR, Bruce T, Zamanian RT, Del Rosario P, Rabinovitch M, et al. Single-Cell Imaging Maps Inflammatory Cell Subsets to Pulmonary Arterial Hypertension Vasculopathy. Am J Respir Crit Care Med. 2024;209:206–218. doi: 10.1164/rccm.202209-1761OC

44. Zhang MQ, Wang CC, Pang XB, Shi JZ, Li HR, Xie XM, Wang Z, Zhang HD, Zhou YF, Chen JW, et al. Role of macrophages in pulmonary arterial hypertension. Front Immunol. 2023;14:1152881. doi: 10.3389/fimmu.2023.1152881

45. Lawther AJ, Phillips AJK, Chung NC, Chang A, Ziegler AI, Debs S, Sloan EK, Walker AK. Disrupting circadian rhythms promotes cancer-induced inflammation in mice. Brain Behav Immun Health. 2022;21:100428. doi: 10.1016/j.bbih.2022.100428

46. Su K, Din ZU, Cui B, Peng F, Zhou Y, Wang C, Zhang X, Lu J, Luo H, He B, et al. A broken circadian clock: The emerging neuro-immune link connecting depression to cancer. Brain Behav Immun Health. 2022;26:100533. doi: 10.1016/j.bbih.2022.100533

47. Shafi AA, Knudsen KE. Cancer and the Circadian Clock. Cancer Res. 2019;79:3806-3814. doi: 10.1158/0008-5472.Can-19-0566

48. Qu M, Zhang G, Qu H, Vu A, Wu R, Tsukamoto H, Jia Z, Huang W, Lenz HJ, Rich JN, et al. Circadian regulator BMAL1::CLOCK promotes cell proliferation in hepatocellular carcinoma by controlling apoptosis and cell cycle. Proc Natl Acad Sci U S A. 2023;120:e2214829120. doi: 10.1073/pnas.2214829120

49. Farshadi E, Yan J, Leclere P, Goldbeter A, Chaves I, van der Horst GTJ. The positive circadian regulators CLOCK and BMAL1 control G2/M cell cycle transition through Cyclin B1. Cell Cycle. 2019;18:16–33. doi: 10.1080/15384101.2018.1558638

50. Simko F, Bednarova KR, Krajcirovicova K, Hrenak J, Celec P, Kamodyova N, Gajdosechova L, Zorad S, Adamcova M. Melatonin reduces cardiac remodeling and improves survival in rats with isoproterenol-induced heart failure. J Pineal Res. 2014;57:177–184. doi: 10.1111/jpi.12154

51. Randhawa PK, Gupta MK. Melatonin as a protective agent in cardiac ischemia-reperfusion injury: Vision/Illusion? Eur J Pharmacol. 2020;885:173506. doi: 10.1016/j.ejphar.2020.173506

52. Tobeiha M, Jafari A, Fadaei S, Mirazimi SMA, Dashti F, Amiri A, Khan H, Asemi Z, Reiter RJ, Hamblin MR, et al. Evidence for the Benefits of Melatonin in Cardiovascular Disease. Front Cardiovasc Med. 2022;9:888319. doi: 10.3389/fcvm.2022.888319

53. Wang R, Zhou S, Wu P, Li M, Ding X, Sun L, Xu X, Zhou X, Zhou L, Cao C, et al. Identifying Involvement of H19-miR-675-3p-IGF1R and H19-miR-200a-PDCD4 in Treating Pulmonary Hypertension with Melatonin. Mol Ther Nucleic Acids. 2018;13:44–54. doi: 10.1016/j.omtn.2018.08.015

54. Chen S, Sun P, Li Y, Shen W, Wang C, Zhao P, Cui H, Xue JY, Du GQ. Melatonin activates the Mst1-Nrf2 signaling to alleviate cardiac hypertrophy in pulmonary arterial hypertension. Eur J Pharmacol. 2022;933:175262. doi: 10.1016/j.ejphar.2022.175262

55. Astorga CR, González-Candia A, Candia AA, Figueroa EG, Cañas D, Ebensperger G, Reyes RV, Llanos AJ, Herrera EA. Melatonin Decreases Pulmonary Vascular Remodeling and Oxygen Sensitivity in Pulmonary Hypertensive Newborn Lambs. Front Physiol. 2018;9:185. doi: 10.3389/fphys.2018.00185

56. de la Fuente A, Zagolín M, Parra V, Paz AA, González-Candia A, Cabrera O, Olave C, Bahamondes C, Gaete MJ, Gudenschwager L, et al. Melatonin Improves Quality of Life, Oxidative Stress, and Cardiovascular Function in Pulmonary Arterial Hypertension. Pulm Circ. 2025;15:e70109. doi: 10.1002/pul2.70109

## References

1. Wang LA, Kern R, Yu E, Choi S, Pan JQ. IntelliSleepScorer, a software package with a graphic user interface for automated sleep stage scoring in mice based on a light gradient boosting machine algorithm. Sci Rep. 2023;13:4275. doi: 10.1038/s41598-023-31288-2

